# Dynamic Control of Argonautes by a Rapidly Evolving Immunological Switch

**DOI:** 10.1101/2024.11.30.626165

**Authors:** Chee Kiang Ewe, Guy Teichman, Maximilian M L Knott, Sarit Anava, Hila Gingold, Mario Bardan Sarmiento, Emily Troemel, Oded Rechavi

**Affiliations:** School of Neurobiology, Biochemistry and Biophysics, Wise Faculty of Life Sciences & Sagol School of Neuroscience, Tel Aviv University, Tel Aviv, Israel; Department of Cell and Developmental Biology, University of California San Diego, California, U.S.A

**Keywords:** RNAi, transgenerational epigenetic inheritance, transgenerational small RNA, Argonuate, innate immunity, antiviral response

## Abstract

Small RNAs, coupled with Argonuate proteins (AGOs), regulate diverse biological processes, including immunity against nucleic acid parasites. *C. elegans* possesses an expanded repertoire of at least 19 AGOs functioning in an intricate gene regulatory network. Despite their crucial roles, little is known about the regulation of AGOs, and whether their expression levels, tissue specificity, and functions change in response to genetic perturbations or environmental triggers. Here, we report that PALS-22, a member of an unusually expanded protein family in *C. elegans*, acts as a negative regulator of antiviral RNAi involving the RIG-I homolog. The loss of *pals-22* leads to enhanced silencing of transgenes and endogenous dsRNAs. We found that PALS-22 normally suppresses the expression of two AGOs, VSRA-1 and SAGO-2, which are activated by bZIP transcription factor ZIP-1. When *pals-22* is eliminated, *vsra-1* and *sago-2* are upregulated. These AGOs in turn play key roles in defense against foreign genetic elements and intracellular pathogens, respectively. Surprisingly, while in *pals-22* mutants immune genes functioning in the intracellular pathogen response (IPR) are upregulated, removing SAGO-2 or the RNA-dependent RNA polymerase RRF-3 in these mutants leads to the downregulation of these genes. This observation contrasts with the typical gene-silencing role of siRNAs. Finally, by analyzing *C. elegans* wild isolates and lab reference strains, we demonstrate that PALS-22 regulates the expression of several germline AGOs, affecting germline mortality and transgenerational epigenetic inheritance. In summary, PALS-22 is a key genetic node that balances the trade-off between immunity and germline health by modulating the functions of different AGOs, thereby shaping the outputs of the RNAi machinery and the dynamics of epigenetic inheritance.

## Introduction

Small RNAs play a pivotal role in gene regulation and influence most, if not all, aspects of animal biology. The central effectors of small RNAs are AGOs characterized by the PAZ and PIWI domains [1–5]. *C. elegans* contains a highly expanded RNAi compared to many other animals, featuring at least 19 functional AGO proteins: 7 in the AGO clade, 1 in the PIWI clade and 11 in the <u>W</u>orm-specific <u>A</u>r<u>go</u>nautes (WAGO) clade, many of which are only partially characterized [1,2,6]. AGOs engage different classes of small RNA including miRNAs, piRNAs and siRNAs. Once loaded onto an AGO, forming RNA-induced silencing complex (RISC), small RNAs guide the complex to sequence-complimentary RNAs targeting them for silencing [3].

AGOs exhibit distinct target repertoire, and in many cases, it is not clear how their specificity in loading particular small RNAs is determined. Among their many roles, the WAGO (or secondary AGOs) family is instrumental in regulating repetitive and transposable elements [1]. Primary AGOs, including RDE-1, ERGO-1, and PRG-1, associate with exogenous siRNAs, endogenous 22G small RNAs, and 21U piRNAs, respectively. Upon target recognition, they trigger the production of abundant amplified 22G “secondary” small RNAs by the RNA-dependent RNA polymerase (RdRP) RRF-1. The 22G small RNAs generated by RRF-1 are loaded onto WAGOs, such as HRDE-1, PPW-1, PPW-2, and WAGO-1, that direct silencing of foreign RNA and also endogenous genes. Unusually, while most AGO-bound small RNAs silence the expression of their targets, 22G RNA produced by RdRP EGO-1 are loaded onto CSR-1, which can instead “license” gene expression in the germline. It is currently unclear how EGO-1 targets are selected, although its subcellular localization to specialized germ granule compartment may play a role [7].

Prior studies demonstrated that CSR-1-associated small RNAs promote transactivation of silenced transgene [8,9] and prevent heterochromatin formation at its target loci [10]. In addition, in sperm, CSR-1 functions with two other AGOs, ALG-3 and −4, to upregulate spermatogenesis genes [11,12]. Interestingly, unlike many other AGOs in *C. elegans*, CSR-1 possesses a slicer domain and can cleave RNA targets. Several studies have shown that it can suppress the expression of embryonic genes in oocytes and tune the levels of maternal mRNA transcripts to ensure proper embryonic development [13,14]. How does CSR-1 balance gene activation and silencing remains unclear. It is important to note that the *csr-1* gene encodes two isoforms that may perform distinct functions in different tissues and developmental stages [12,15].

In many animals, loss of PIWI and AGO leads to infertility and lethality in part due to elevated transposon mobilization and chromosomal segregation defects in the gametes and developing embryos [16]. In *C. elegans*, CSR-1 is required for proper chromosomal pairing, segregation, and early embryonic development [13,14,17,18]. Animals lacking *csr-1* are nearly sterile and produce few arrested embryos [18]. On the other hand, mutants lacking the piRNA pathway or WAGOs do not show overt developmental defects but exhibit progressive sterility in the presence of heat stress (termed mortal germline “Mrt” phenotype) owing to transgenerational accumulation of epigenetic baggage and dysregulation of histone genes, rRNA, and transposable elements [19–21]. Interestingly, a recent study demonstrated that many *C. elegans* wild isolates showed natural variation in Mrt phenotype, which can be rescued by associating the animals with their natural microbiome and pathogens, highlighting the role of environmental factors in the regulation of transgenerational epigenetic inheritance mechanisms [22] – a finding highly relevant to the results presented in this study (see below). While the mechanism of genomic surveillance is extensively studied in the germline, the regulation of non-self genes in the soma is less well understood. In the soma, the regulation of repetitive and transposable elements is far more relaxed than in the germline. The elevated transposon mobilization in the soma was suggested to contribute to phenotypic plasticity and cellular heterogeneity, particularly in the nervous system [23–26]. On the other hand, uncontrolled somatic transposition has been linked to various human diseases such as psychiatric disorders [27,28], neurodegenerative diseases [29,30], and cancer [31].

Among their many important functions, small RNA pathways play a crucial role in immunity in plants and animals. *C. elegans* lacks NF-κB signaling and canonical adaptive immunity; however, it contains homologs of RIG-I-like receptor (RLR) which are conserved RNA sensors that detect viral infection [32]. Together with the Dicer protein DCR-1 and the dsRNA binding protein RDE-4, DRH-1/RIG-I forms an antiviral complex which processes viral RNAs into 23 nucleotides long primary “viRNAs”. These viRNAs are then loaded onto the primary AGO RDE-1, which recruits RRF-1 to the viral target transcripts to generate secondary siRNAs, which are subsequently loaded onto WAGOs, such as SAGO-2, to promote degradation of complementary viral RNA targets (**Figure 1A**) [1,33–36].

**Figure 1:**
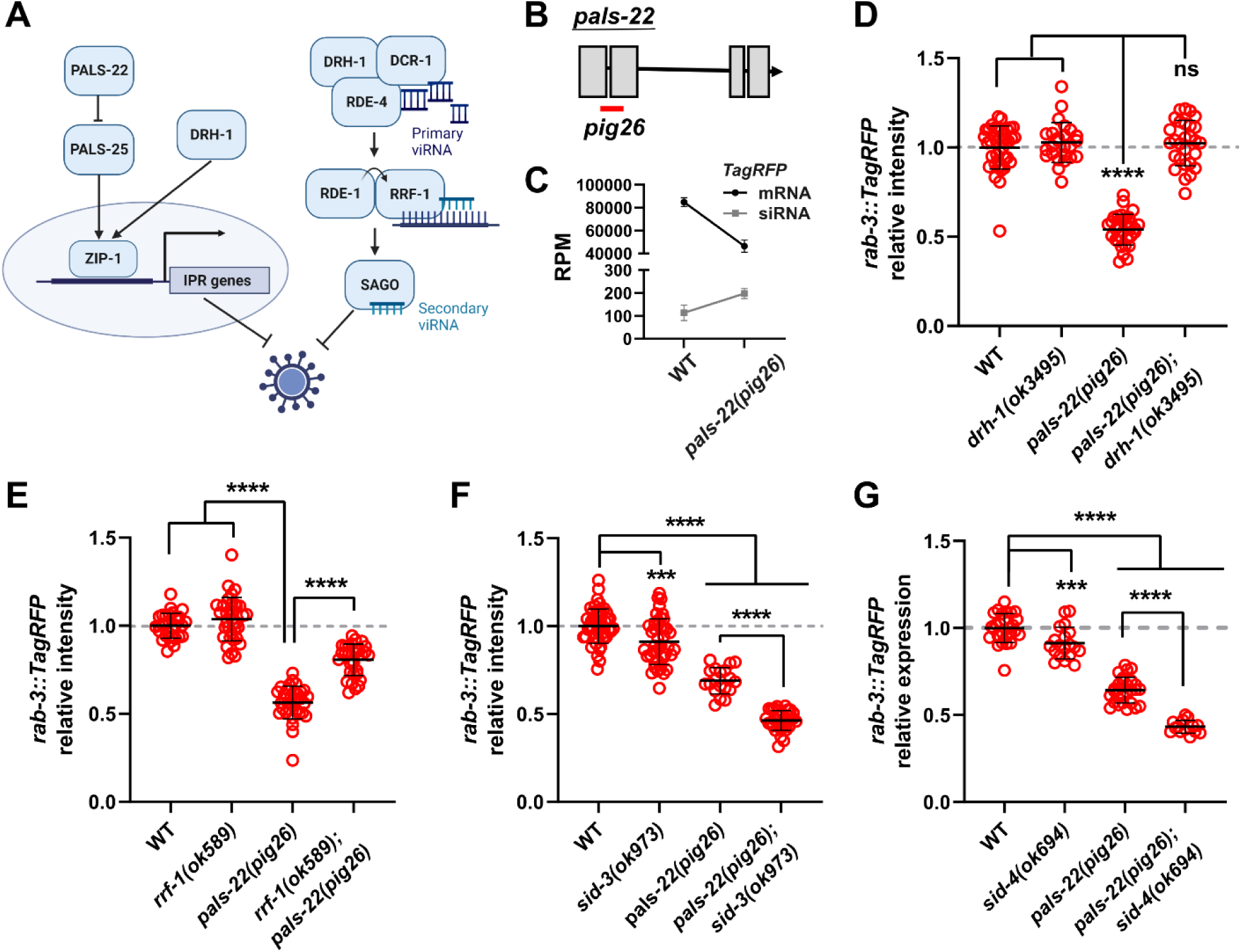
The antiviral RNAi pathway and conserved kinase signaling regulate the expression of repetitive transgenes. (A) Intracellular pathogen response and RNAi defense against viral pathogen in *C. elegans*. (B) *pig26* allele contains a 253 bp deletion in *pals-22* indicated by the red bar. (C) mRNA and antisense siRNA reads mapped to *rab-3p::TagRFP* transgene in wild-type and *pals-22(pig26)* animals. RPM: reads per million. (D, E) Requirement of DRH-1 and RRF-1 for transgene silencing in *pals-22(pig26)*. (F, G) Interactions between SID-3, −4 and PALS-22 in transgene silencing. Each data point represents an animal. Statistical significance was determined by parametric one-way ANOVA followed by pairwise two-tailed unpaired t-tests with Benjamini–Hochberg correction. ns, not significant (q > 0.05); *** q ≤ 0.001; **** q ≤ 0.0001. Error represents mean +/− SD.

Aside from using RNAi to target viruses, it was recently found that *C. elegans* mounts a transcriptional response against intracellular pathogens such as Orsay virus (the sole known natural virus of *C. elegans*) and microsporidia *N. parisii,* which is a fungal pathogen [34,35,37–42]. This intracellular pathogen response (IPR) is in part governed by an unusually expanded *pals* (short for <u>p</u>rotein containing <u>ALS</u>2CR12 signature) gene family. It is noteworthy that *C. elegans* contains 39 copies of *pals* genes, compared to 8 in its cousin *C. briggsae* [43]. This rapid proliferation of protein coding genes is extraordinary and may reflect intense host-pathogen arms race that drives escalated diversification of innate immunity.

One of the members of the *pals* family, *pals-22*, is known to function as a repressor of the IPR by inhibiting *pals-25*. Loss of *pals-22* in worms enhances their resistance to viral and microsporidian pathogens in *pals-25*-dependent manner [39]. Interestingly, DRH-1 may also activate IPR gene expression independently of RNAi factors, much like mammalian RLR [44,45]. Upon infection, PALS-25 and DRH-1 induce upregulation and nuclear localization of the ZIP- 1/bZIP transcription factor, triggering IPR gene expression and systemic antiviral defense (**Figure 1A**) [32,35,46]. Curiously, eliminating *pals-22* causes enhanced silencing of repetitive transgenes and enhanced exogenous RNAi response (classical “Eri” phenotypes) [43], poising PALS-22 at a critical node that regulates a broad range of genomic parasites, although the mechanism of action of PALS-22 in the RNAi pathways remains elusive.

In this study, we sought to investigate the mechanism by which PALS-22 regulates RNAi. This investigation uncovered the role of an unusual WAGO, VSRA-1 (<u>V</u>ersatile <u>S</u>mall <u>R</u>NAs <u>A</u>rgonaute-1; also named CSR-2 [17]), in regulating repetitive foreign elements in somatic tissues. Our analyses further revealed that PALS-22 controls the levels, expression pattern, and activities of multiple AGOs, which in turn regulate the expression of transgenes and immune genes. Interestingly, we found that the small RNA system assumes two distinct states, depending on the activity of PALS-22, and exerting opposing effects on IPR gene expression. Specifically, the endo-siRNA pathway, which involves SAGO-2, normally silences IPR genes; however, in the absence of PALS-22, it instead upregulates their expression. Moreover, we report that natural variation in the *pals-22* gene strongly affects the Mrt phenotype, linked to transgenerational epigenetic inheritance mechanisms. Together, our work demonstrates that PALS-22 controls the gene regulatory outputs of multiple AGOs and further uncovers a potential cryptic role of SAGO-2 in supporting gene expression. This rapidly evolving genetic module may play a key role in how the animals interact with pathogens in their natural environments, potentially influencing epigenetic inheritance and evolutionary trajectory of the animals.

## Results

### Loss of *pals-22* enhances repetitive transgene silencing via the antiviral RNAi pathway

To study the roles of PALS-22 in the RNAi pathways, we used CRISPR-Cas9 to generate a deletion allele of *pals-22* (designated *pig26*) (**Figure 1B**). We found that *pals-22(pig26)* exhibits many phenotypes of previously characterized loss-of-function mutations of *pals-22*, including increased repetitive transgene silencing [43] (**Figure 1C**; **Figure S1A, B)**, enhanced resistance to Orsay virus and *N. parisii* infection [39] (see below), decreased brood size (**Figure S1C**), and age-dependent locomotory defects (**Figure S1D**) [43].

LIN-35/Rb and the DREAM/Muv B complex repress germ cell fate in the soma. Inactivating *lin-35* leads to a partial soma-to-germline transformation, as evidenced by the presence of ectopic germline P granules, and enhanced RNAi [47]. Intriguingly, a recent study demonstrated that the loss of *lin-35/Rb* induced a transcriptional response similar to that of *pals-22* mutants and *N. parisii* -infected worms [48]. Therefore, we asked whether the enhanced transgene-silencing activity in *pals-22(pig26)* is the result of de-repression of germline RNAi in somatic cells, as observed in *lin- 35(-)* mutants. However, we did not observe obvious misexpression of the germline P granule component PGL-1 in the soma of *pals-22(pig26)* mutants (**Figure S2**).

The nuclear RNAi factor NRDE-3 directs deposition of H3K9 methylation and induces transcriptional silencing of repetitive foreign elements [49]. However, we found that enhanced transgene silencing in *pals-22(pig26)* does not depend on NRDE-3. While NRDE-3 localizes to the cytoplasm in many “Eri” mutants due to absence of endo-siRNAs [50], it remains prominently nuclear in *pals-22* mutants (**Figure S3A**). Additionally, knocking out *nrde-3* or *set-25/HMT* does not affect *pals-22(pig26)* transgene-silencing phenotype (**Figure S3B**). On the other hand, knocking out *drh-1* or *rrf-1* strongly suppresses the silencing phenotype of *pals-22(pig26)* mutants (**Figure 1D, E**), suggesting that the loss of *pals-22* triggers post-transcriptional cytoplasmic silencing of repetitive element by the antiviral RNAi pathway. Concordant with RNAi-mediated silencing of transgenes, we observed a significant upregulation of siRNAs targeting *TagRFP* in *pals-22(pig26)* mutants (log_2_FC = 0.70, DeSeq2 q = 0.02; **Figure 1C**). In line with the hyperactive RNAi phenotype following the loss of *pals-22*, genes associated with the GO term “regulatory ncRNA-mediated post-transcriptional gene silencing” are significantly enriched among those with increased expression in *pals-22(pig26)* mutants versus wild-type worms (GO: 0035194; 22 observed and 8.36 expected; Fisher’s exact test, q = 0.005) (**Figure S4A**). Indeed, *drh-1*, along with genes encoding core components of the RNAi pathway, such as *rde-1*, *rde-4*, and *rrf-1*, are upregulated in *pals-22(pig26)* mutants (**Figure S4B**). Additional genes encoding factors involved in the amplification of secondary siRNAs, including *ekl-1*, *drh-3*, *rsd-2*, and *rde-12*, also showed elevated expression in *pals-22* mutants relative to the wild-type worms (**Figure S4B**).

Prior studies reported that *sid-3/TNK2*, which encodes a cytoplasmic tyrosine kinase implicated in RNA import and endosomal trafficking, is critical for Orsay virus infection [38,51,52]. Eliminating SID-3 enhances resistance to viral infection in worms [38]. Interestingly, we found that knocking out *sid-3* leads to a modest but statistically significant drop in transgene expression (q = 0.0004; **Figure 1F**). Moreover, *pals-22(pig25); sid-3(ok973*) double mutants show stronger silencing than either of the single mutants (**Figure 1F**). In humans, TNK2 is known to interact with an adaptor protein NCK1 to promote virus entry and facilitate subsequent proper endocytic trafficking of the viral particles [38,53]. Consistently, knocking out *sid-4/NCK1* (also named *nck-1*) enhances *pals-22’s* transgene-silencing phenotype (**Figure 1G**). Together, our results demonstrate that antiviral immunity and repetitive DNA are co-regulated by PALS-22 and a conserved kinase signaling pathway. Further investigation is warranted to elucidate the mechanism by which SID-3 and SID-4 promote the expression of repetitive foreign genetic elements.

### PALS-22 represses expression of the Argonaute protein VSRA-1

The foregoing results suggest that the antiviral RNAi response promotes transgene silencing in the soma. This led us to test the functions of the SAGO-2/WAGO, which has previously been shown to be important for fighting viral infections [1,33]. However, we found that removing *sago- 2* either by RNAi or a null mutation (*tm894*) does not alter the expression of repetitive transgenes in both wild type and *pals-22(pig26)* mutants (**Figure 2A; Figure S5**), suggesting that the antiviral RNAi pathway engages a different WAGO to distinguish exogenous/virus-derived RNA from repetitive transgenes.

**Figure 2:**
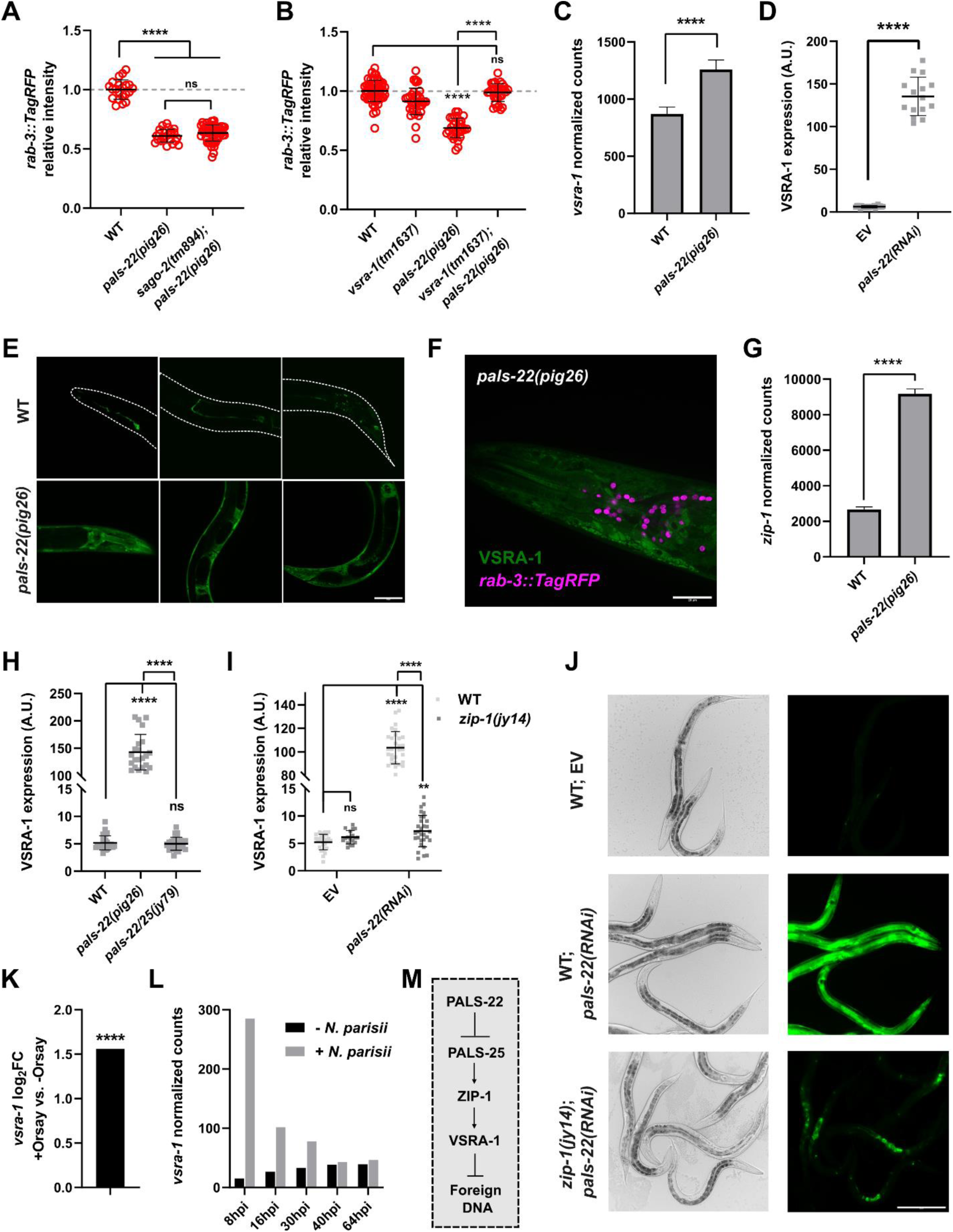
PALS-22 represses VSRA-1. (A) Requirement of SAGO-2 for transgene silencing in *pals-22(pig26)*. (B) Requirement of VSRA- 1 for transgene silencing in *pals-22(pig26)*. Statistical significance was determined by parametric one-way ANOVA followed by pairwise two-tailed unpaired t-tests with Benjamini–Hochberg correction. ns, not significant (p > 0.05); *** q ≤ 0.001; **** q ≤ 0.0001. Error represents mean +/− SD. (C) Expression of *vsra-1* in wild type and *pals-22(pig26).* The mRNA read counts were normalized with relative log expression normalization method. **** DESeq2 q ≤ 0.0001. (D) Expression of endogenous VSRA-1 reporter in wild-type and *pals-22(RNAi)* animals. Day 1 adults were scored. Scale bar = 100 µm. A.U. = arbitrary unit. Statistical significance was determined by non-parametric Mann-Whitney test. **** p ≤ 0.0001. Error represents mean +/− SD. (E) Tissue localization of VSRA-1 in wild-type and *pals-22(pig26)* day 1 adults. Scale bar = 50 µm. (F) Accumulation of VSRA-1 in the cytoplasm in *pals-22(pig26)*. Neuronal nuclei are marked by *otIs355[rab-3p::TagRFP]*. Scale bar = 20 µm. (G) Expression of *zip-1* in wild type and *pals- 22(pig26).* The mRNA read counts were normalized with relative log expression normalization method. **** DESeq2 q ≤ 0.0001 (H) Expression of VSRA-1 endogenous reporter in *pals- 22(pig26)* compared to *pals-22/25(jy79)* animals. (I, J) Expression of VSRA-1 endogenous reporter in *pals-22(RNAi)* compared to *zip-1(jy14); pals-22(RNAi)* animals. Statistical significance was determined by non-parametric Kruskal-Wallis test followed by pairwise Mann-Whitney tests with Benjamini–Hochberg correction. ns, not significant (q > 0.05); **** q ≤ 0.0001. Error represents mean +/− SD. Scale bar = 200 µm. (K) Expression of *vsra-1* during infection with Orsay virus. **** q ≤ 0.0001. The data was extracted from Chen, *et. al.* [56] (L) Expression of *vsra-1* during infection with *N. parasii*. The data was extracted from Bakowski, *et. al.* [37]. hpi = hours post inoculation. The mRNA read counts were normalized with FPKM method. (M) A model indicating the regulation and function of VSRA-1.

By examining Co-IP mass spectrometry data of PALS-22 [54], we noted that PALS-22 physically interacts with VSRA-1, a close paralog of CSR-1 [1,6]. We found that deleting *vsra-1* strongly rescues transgene silencing in *pals-22(pig26)* (**Figure 2B; Figure S6**). We found that *vsra-1* transcripts are highly upregulated in *pals-22(pig26)* (**Figure 2C**). Consistently, by examining an endogenous fluorescent translational reporter of VSRA-1 (CRISPR-mediated tagging of the protein), we found that VSRA-1 is dramatically upregulated in *pals-22(pig26*) and *pals-22(RNAi*) animals (**Figure 2D-F**). This upregulation is observed across all developmental stages (**Figure S7A**). Moreover, while VSRA-1 is only detectable in the somatic gonad and a small number of neurons in wild-type animals (**Figure 2E; Table S1**), we found that it is expressed ubiquitously in *pals-22(pig26)* mutants (**Figure 2E**). VSRA-1 appears to exclusively localize to the cytoplasm (**Figure 2F**), suggesting that VSRA-1 functions in post-transcriptional gene regulation.

ZIP-1/bZIP is a major transcription factor inducing IPR; ZIP-1 is activated by PALS-25 and eliminating *zip-1* leads to increased susceptibility to intracellular pathogens (see **Figure 1A**) [46,55]. We found that *zip-1* is significantly upregulated in *pals-22(pig26)* mutants in which the IPR is constitutively activated (**Figure 2G**), in agreement with previous studies [46,55]. Hence, we hypothesized that *vsra-1* may be activated by ZIP-1 downstream of PALS-25. Indeed, deleting *pals-25* suppresses *vsra-1* expression in *pals-22(pig26)* mutants (**Figure 2H**). Furthermore, knocking down *zip-1* by RNAi significantly downregulates *vsra-1* in *pals-22(pig26)* mutant background (**Figure S7B, C**). We observed a similar effect when *pals-22* is eliminated by RNAi in *zip-1(jy14)* mutants (**Figure 2I**). However, intriguingly, *vsra-1* remains upregulated in the somatic gonad of *zip-1(jy14); pals-22(RNAi)* animals (**Figure 2J**), suggesting PALS-22 engages tissue-specific gene regulatory modules. The biological significance of elevated *vsra-1* expression in the somatic gonad is currently unclear, but it is worth noting that it is the only tissue, besides the intestine, that is susceptible to Orsay virus infection [42].

Interestingly, by examining published transcriptomic data, we found that *vsra-1* is upregulated in worms infected with Orsay virus (**Figure 2K**) [56] or *N. parisii* (**Figure 2L**) [37]. During *N. parisii* infection, this upregulation is especially pronounced at early time-points that correspond to the entry and replication of the parasite inside host intestinal cells (**Figure 2L**) [37,41]. On the contrary, *vsra-1* expression remains unchanged in animals exposed to extracellular bacterial and other fungal pathogens [57]. Taken together, these results indicate that VSRA-1 responds to the presence of intracellular pathogens, downstream of the PALS-25◊ZIP- 1 regulatory axis, and mounts an RNAi defense to mitigate the infection (**Figure 2M**). Supporting this notion, knocking down *vsra-1* by RNAi causes increased susceptibility to viral infection [58].

### PALS-22 promotes the expression of endogenous dsRNAs and silences newly evolved genes

To further probe the roles of PALS-22 in somatic small RNA pathways, we sequenced small RNAs and mRNAs isolated from wild-type and *pals-22(pig26)* L1 animals. The small RNAs were cloned using a 5′-phosphate-independent protocol to capture both primary and secondary small RNAs (see **Methods**). We chose to sequence L1 animals because they contain 550 somatic cells and only two primordial germ cells, and thus any transcriptional changes are likely to be indicative of somatic responses. Consistent with our previous findings that the antiviral RNAi pathway is enhanced in the absence of *pals-*22, we found that the overall levels of 22G secondary siRNAs are significantly elevated in *pals-22(pig26)* (**Figure 3A, B**).

**Figure 3:**
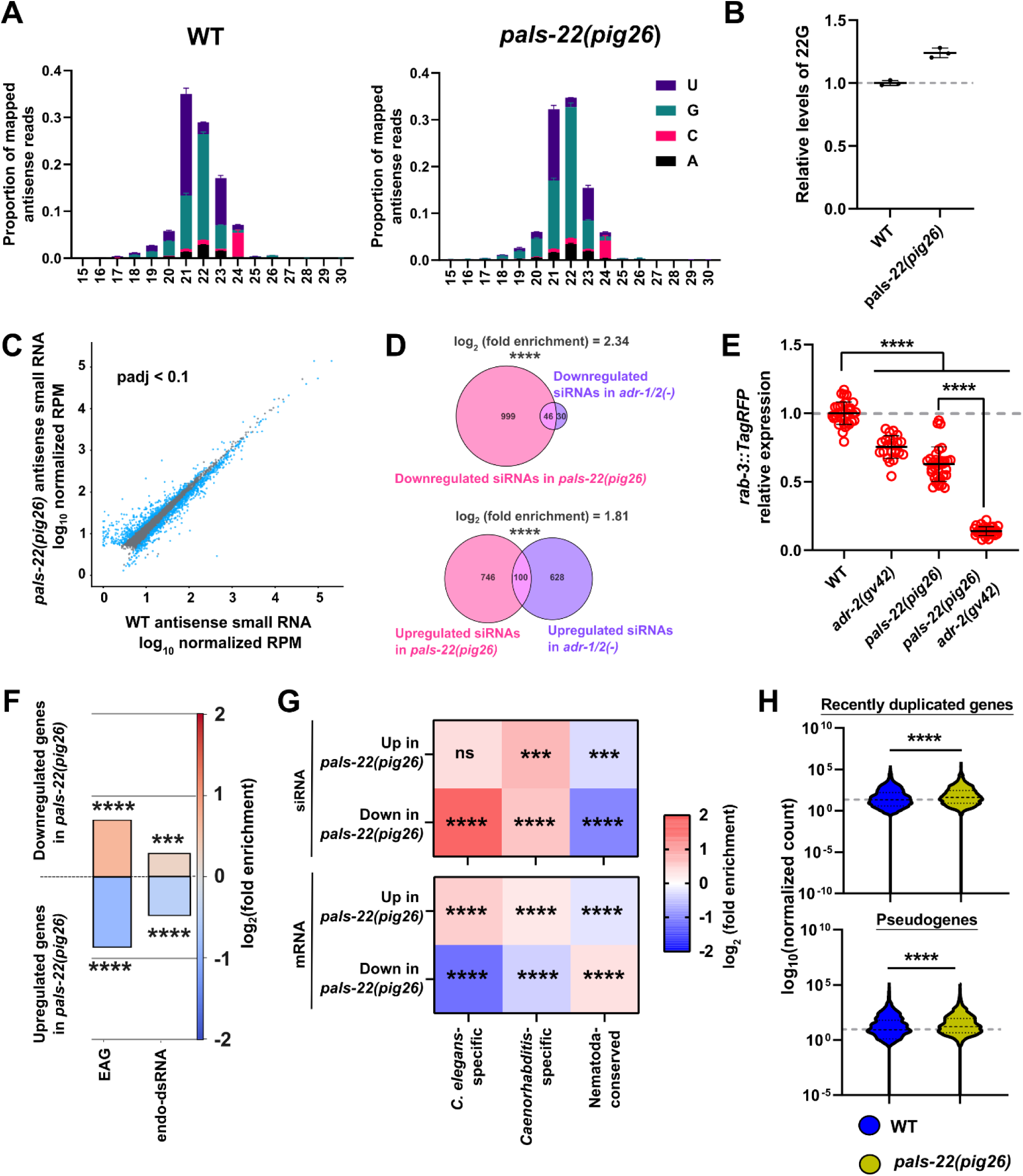
Loss of *pals-22* leads to remodeling of the small RNA pool. (A) Antisense small RNA size and 5’ nucleotide distribution in wild type and *pals-22(pig26)*. (B) Levels of 22G small RNA in *pals-22(pig26)* normalized to wild type. (C) Differentially expressed siRNAs (DESeq2 padj < 0.1) in *pals-22(pig26)* compared to wild type. RPM: read per million. (D) Overlaps of differentially expressed siRNAs (DESeq2 padj < 0.1) in *pals-22(pig26)* and *adr-1/2(-)*. **** q ≤ 0.0001. (E) Interactions between PALS-22 and ADR-2 in transgene silencing. Statistical significance was determined by non-parametric Kruskal–Wallis test followed by pairwise Wilcoxon Rank Sum tests with Benjamini–Hochberg correction. **** q ≤ 0.0001. Error represents mean +/− SD. (F) Enrichment analysis for A-to-I editing-associated genes (EAG) and endo-dsRNA (ΔG/nt < −0.5 kcal/mol × nt) in *pals-22(pig26)* differentially expressed mRNAs. *** q ≤ 0.001; **** q ≤ 0.0001. (G) Enrichment analysis for *C. elegans*-specific, *Caenorhabditis*-specific, and Nematoda- conserved genes in *pals-22(pig26)* differentially expressed siRNAs and mRNAs. ns, not significant (q > 0.05); *** q ≤ 0.001; **** q ≤ 0.0001. (H) Expression of recently duplicated genes and pseudogenes in wild type and *pals-22(pig26)*. The mRNA read counts were normalized with relative log expression normalization method. Statistical significance was determined by non- parametric Mann-Whitney test. **** p ≤ 0.0001.

Loss of *pals-22* causes differential expression of many siRNAs targeting protein-coding and non-protein coding genes (DESeq2 padj < 0.1; **Figure 3C**). By taking a closer examination at the gene biotypes, we found that siRNAs targeting protein-coding genes, lincRNAs, and pseudogenes are upregulated in *pals-22(pig26)*, while the levels of miRNAs and piRNAs remain largely unchanged (**Figure S8**).

In *C. elegans*, zebrafish, and mammals, ADARs are required to prevent ectopic activation of the antiviral response, involving RLR, targeting self dsRNAs [59–61]. In vertebrate, recognition of dsRNAs by MDA-5 and RIG-I triggers IFN-response [60,61]. In *C. elegans*, RNA editing disrupts dsRNA formation and thus antagonizes the production of dsRNA-derived siRNAs by DCR-1 [60,62,63]. Loss of ADARs, like PALS-22, leads to transgene silencing and reduced lifespan (see **Figure S1**) [43,64]. Hence, we asked whether PALS-22 may interact with ADARs to regulate the expression of transgene-derived and endogenous dsRNAs. Supporting our hypothesis, we found that the small RNA profile of *pals-22(pig26)* significantly overlaps with that of *adr-1/2(-)* mutants (**Figure 3D**). Indeed, we found that PALS-22 and ADR-2 function synergistically to promote repetitive transgene expression as *pals-22(pig26) adr-2(gv42)* double mutant shows a stronger silencing phenotype than either of the single mutants (**Figure 3E**). Moreover, many A-to-I editing-associated genes (EAG) previously defined by Reich *et al.* [62] and endo-dsRNA containing structured introns (ΔG/nt < −0.5 kcal/mol × nt) [65] are downregulated in *pals-22(pig26)* (**Figure 3F**). These findings suggest that PALS-22 prevents aberrant silencing of endogenous dsRNAs likely by suppressing the antiviral RNAi pathway.

Evolutionarily young genes tend to be the targets of endogenous siRNA pathways [66–68]. On the other hand, dsRNA-associated genes targeted by ADAR are enriched among conserved, essential genes [65,69]. We found that the loss of *pals-22* causes downregulation of siRNAs targeting newly evolved genes, such as *C. elegans*-specific genes and genes that have no orthologs outside of the *Caenorhabditis* genus (**Figure 3G**). Concordantly, we found a significant enrichment for newly evolved genes, including pseudogenes and young duplicated genes, among the mRNAs that are upregulated in *pals-22(pig26)* (**Figure 3G, H**). In contrast, downregulated mRNAs in *pals-22(pig26)* are enriched for older genes that are conserved across the Nematoda phylum (**Figure 3G**). Hence, PALS-22 blocks the expression of young genes while promoting the expression of conserved genes.

Collectively, our findings indicate that the loss of *pals-22* drastically remodels the small RNA landscape and imparts a variety of gene regulatory outcomes on targets’ transcripts with distinct features, possibly including secondary structures, number of introns, and divergent splicing signals, recognized by the RNAi machinery [66,68].

### Dual states of RRF-3-dependent endo-siRNAs in controlling IPR gene expression

In *C. elegans*, small RNAs can be grouped into four distinct clusters based on the AGOs they tend to associate with (**Figure 4A**) [1]. We noted that differentially expressed siRNAs in *pals-22(pig26)* tend to associate with the “WAGO cluster” composed of WAGO-1, PPW-2, HRDE-1, and PPW-1, which are major regulators of repetitive and transposable elements [1] (**Figure 4A**). Furthermore, upregulated siRNAs in *pals-22(pig26)* are enriched for the “ERGO-1 cluster” (composed of SAGO-2, SAGO-1, ERGO-1, and NRDE-3) targets, which are largely associated with immune, defense, and stress responses (**Figure 4A**) [1].

**Figure 4:**
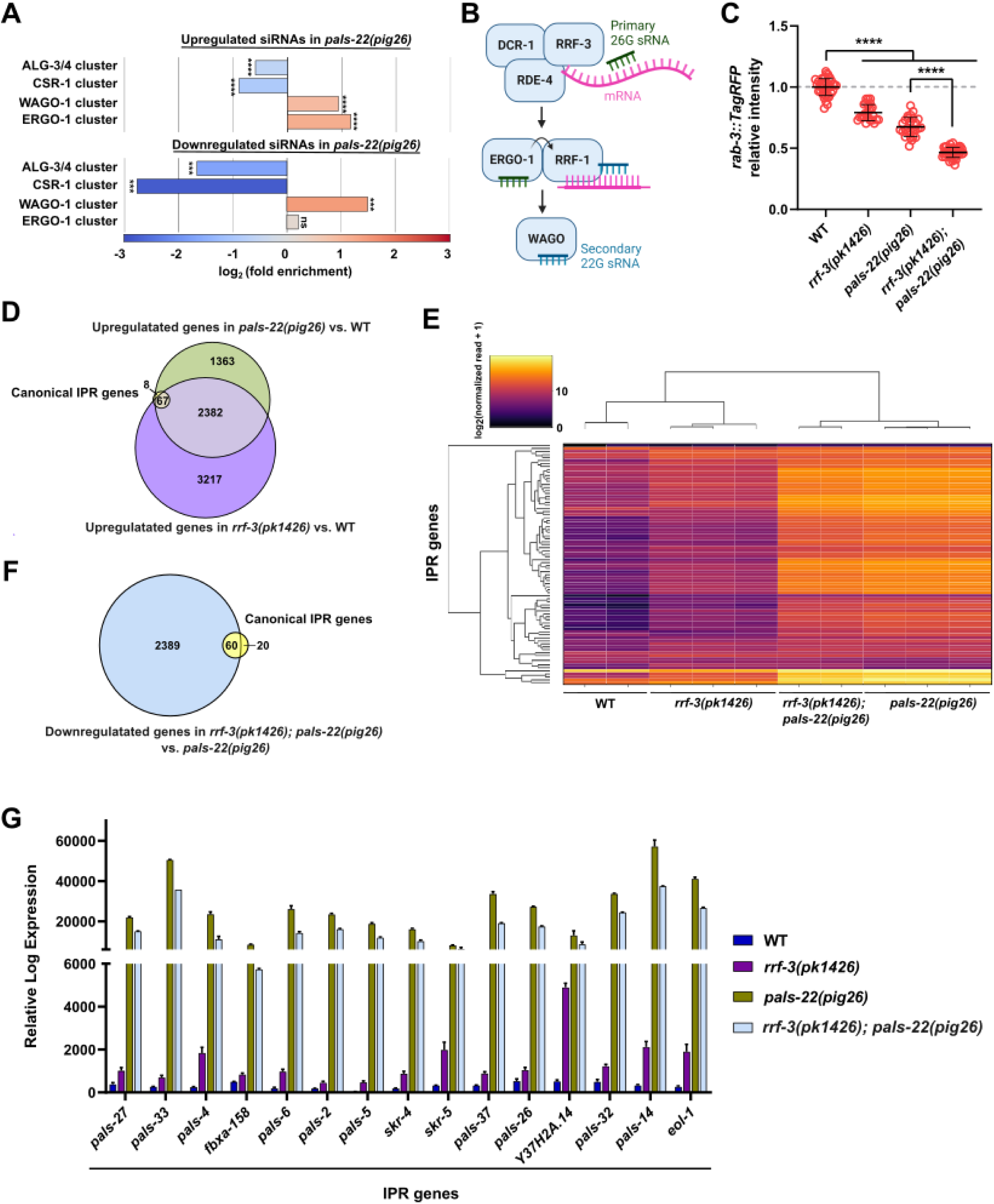
Dual roles of endo-siRNA in regulating IPR gene expression. (A) Enrichment analysis for AGO-associated small RNAs in *pals-22(pig26)* differentially expressed siRNAs. Endo-siRNA-associated AGOs are divided into four clusters [1]. (B) Overview of the 26G endo-siRNA pathway. (C) Interactions between PALS-22 and RRF-2 in transgene silencing. Statistical significance was determined by non-parametric Kruskal–Wallis test followed by pairwise Wilcoxon Rank Sum tests with Benjamini Hochberg correction. **** q ≤ 0.0001. Error represents mean +/− SD. (D) Venn diagram showing the overlap between 80 canonical IPR genes and genes upregulated in *pals-22(pig26)* and *rrf-3(pk1426)* mutants compared to wild type. (E) Hierarchical clustergram plot visualizing the expression of IPR genes in wild type, *rrf-3(pk1426)*, *rrf-3(pk1426); pals-22(pig26)*, and *pals-22(pig26)* using Euclidean distance metric and average linkage. (F) Venn diagram showing the overlap between 80 canonical IPR genes and genes downregulated in *rrf-3(pk1426); pals-22(pig26)* double mutants compared to *pals-22(pig26)* single mutants. (G) Examples of IPR genes upregulated in *rrf-3(pk1426)* compared to wild type but downregulated in *rrf-3(pk1426); pals-22(pig26)* compared to *pals-22(pig26).* Error represents mean +/− SD.

RRF-3 is an RNA-dependent RNA polymerase which functions in the endo-siRNA pathway, upstream of ERGO-1, to generate primary 26G small RNAs using the target mRNA as template (**Figure 4B**) [2]. Consistent with previous findings [70], knocking out *rrf-3* promotes silencing of repetitive transgene (**Figure 4C**). We found that loss of *pals-22* further enhances the *rrf-3* transgene-silencing phenotype (**Figure 4C**), indicating that RRF-3 and PALS-22 function in parallel to antagonize RNAi against foreign genes.

Next, to probe how PALS-22 and the endo-siRNA pathway regulate immunity, we performed RNA-seq analysis and examined IPR gene expression in *rrf-3(-)* and *pals-22(-)* mutants. As expected, eliminating *pals-22* causes a dramatic increase in IPR gene expression compared to wild type (**Figure 4D, E**) [39,71]. Similarly, IPR genes are upregulated in *rrf-3(pk1426)* compared to wild type, albeit to a lesser extent, suggesting that the 26G endo-siRNA pathway normally silences IPR genes (**Figure 4D, E**). As IPR genes are strongly upregulated in *pals- 22(pig26)*, we expected to observe a decrease in the levels of small RNAs that target these genes. To our surprise, we witnessed that exact opposite: small RNAs targeting IPR genes are significantly upregulated in *pals-22(pig26)* mutants (**Figure S9A**) and are positively correlated with the expression of IPR transcripts (Spearman’s ρ = 0.34, slope = 0.47) (**Figure S9B**), in contradiction to the typical gene-silencing role of siRNAs. We sought to further investigate this unusual pattern of gene regulation by examining the interactions between PALS-22, RRF-3, and downstream WAGOs in this process (see below).

Intriguingly, while IPR genes are upregulated in each of the single mutants, 60 out of 80 IPR genes are significantly downregulated in the *rrf-3(pk1436); pals-22(pig26)* double mutant compared to *pals-22(pig26)* (DeSeq2 q < 0.1; **Figure 4E-G**). However, we found that eliminating *rrf-3* in either wild-type or *pals-22(pig26)* animals does not overtly affect their resistance to Orsay virus or *N. parisii* (**Figure S10A-C**).

Together, our results suggest that the small RNA pathway may assume two different states for modulating IPR gene expression: in the wild-type background, the 26G endo-siRNA pathway, acting through RRF-3, represses IPR gene expression; however, removing *pals-22* flips the state such that RRF-3 positively regulates IPR genes in the *pals-22(pig26)* mutant.

### PALS-22 regulates a negative and positive gene expression switch via SAGO-2

As we have shown that VSRA-1 silences transgenes in *pals-22(-)* mutants, we next asked whether VSRA-1 may function downstream of PALS-22 and RRF-3 to regulate IPR gene expression. We found that IPR genes are modestly upregulated in *vsra-1(tm1637)* single mutants; however, unlike *rrf-3*, knocking out *vsra-1* does not alter IPR gene expression in *pals-22(pig26)* mutants (**Figure 5A**).

**Figure 5:**
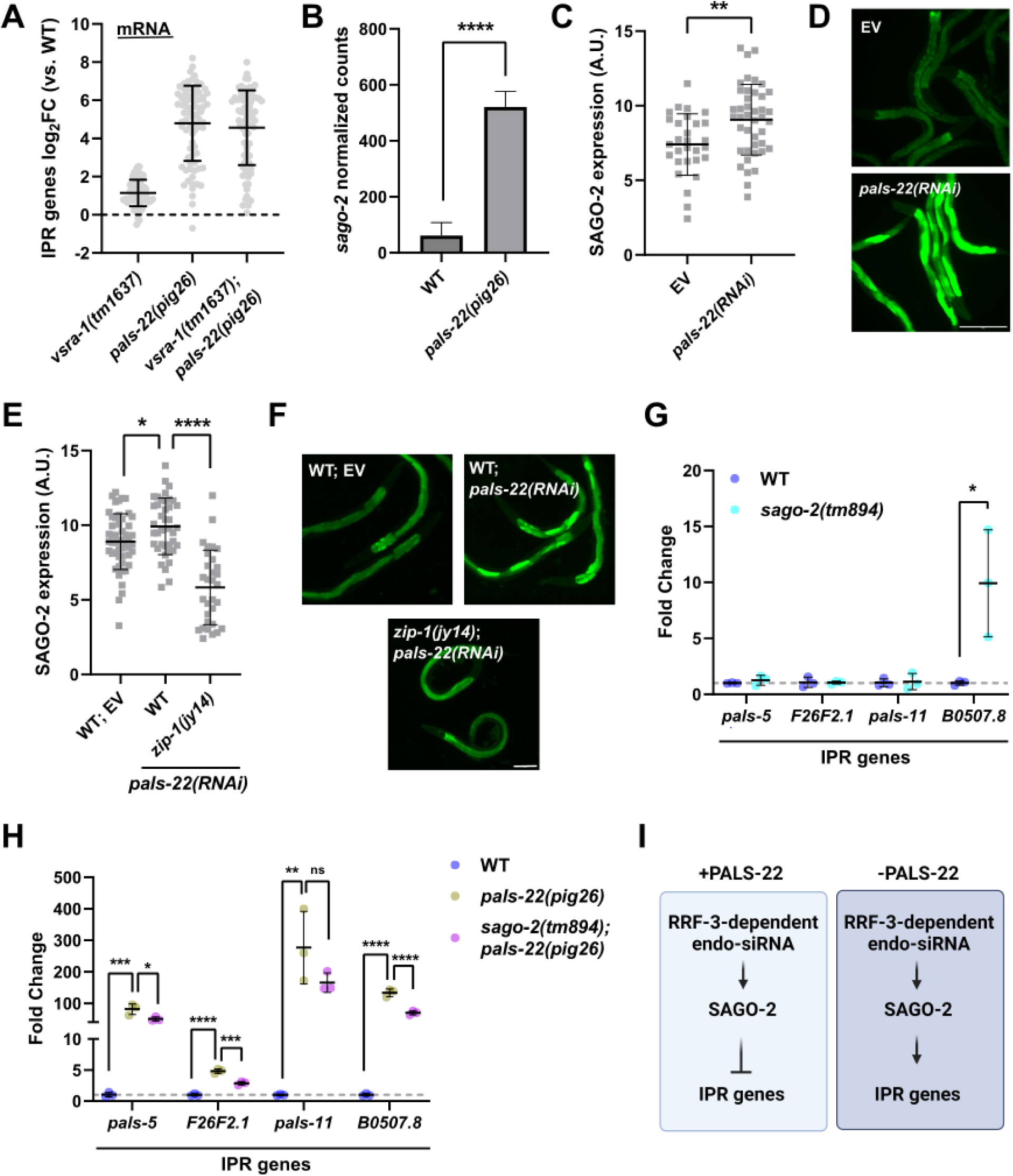
SAGO-2 regulates IPR gene expression. (A) The expression of 80 canonical IPR genes in *vsra-1(tm1637)*, *pals-22(pig26)*, and *vsra- 1(tm1637); pals-22(pig26)*. (B) The expression of *sago-2* transcript in *pals-22(pig26)*. The mRNA read counts were normalized with relative log expression normalization method. **** DESeq2 q ≤ 0.0001. (C, D) The expression SAGO-2 endogenous reporter in animals treated with empty vector (EV; L4440) and *pals-22* RNAi. Statistical significance was determined by parametric two- tailed unpaired t-test. ** p ≤ 0.01. Scale bar = 200 µm. (E-F) The expression SAGO-2 endogenous reporter in wild-type and *pals-22(RNAi)* animals compared to *zip-1(jy14); pals- 22(RNAi)* animals. Statistical significance was determined by parametric one-way ANOVA followed by pairwise two-tailed unpaired t-tests with Benjamini–Hochberg correction. * q < 0.05; **** q ≤ 0.0001. Scale bar = 100 µm. (G) The expression of IPR genes in wild type and *sago- 2(tm894)*, as measured by qPCR. Statistical significance was determined by non-parametric Mann-Whitney test. * p ≤ 0.05. (H) The expression of IPR genes in wild type, *pals-22(pig26)*, *sago-2(tm894); pals-22(pig26)*, as measured by qPCR. All strains contain *otIs355[rab- 3p::TagRFP]* transgene. Statistical significance was determined by non-parametric Kruskal– Wallis test followed by pairwise Wilcoxon Rank Sum tests with Benjamini–Hochberg correction. ns, not significant (q > 0.05); * ≤ 0.05; ** ≤ 0.01; *** q ≤ 0.001; **** q ≤ 0.0001. Error represents mean +/− SD. (I) A model indicating the dual states of endo-siRNA pathway, which inhibits or promotes IPR gene expression, depending on the activity of PALS-22.

In search for a different AGO that could regulate IPR gene expression, we noted that *sago- 2*, like *vsra-1*, is significantly upregulated when *pals-22* is eliminated by a null mutation or RNAi (**Figure 5B-D**). Additionally, we observed that small RNAs associated with SAGO-2 are upregulated in *pals-22(pig26)* mutants (**Figure S11A, B**). Similar to *vsra-1*, we found that PALS- 22 normally represses *sago-2* expression by suppressing the PALS-25→ZIP-1 genetic module. Knocking out *zip-1* in *pals-22(RNAi)* animals restores *sago-2* expression to wild-type levels (**Figure 5E, F**).

To search for additional regulators of *sago-2*, we leveraged the extensive natural variation of *sago-2* expression among *C. elegans* wild isolates (**Figure S12A**) and performed an eQTL analysis on the CaeNDR platform [72–75]. This revealed a local QTL on Chr I and two distant QTLs on Chr II and III (**Figure S12B, C**). We subsequently performed a targeted RNAi screen, focusing on candidates in Chr II, and found that knocking out *cpna-1/F31D5.3* by RNAi in the wild-type background modestly reduces *sago-2* expression (**Figure S12D**). Interestingly, *cpna-1* encodes a Copine Domain protein, which has been implicated in immune responses from plants to mammals [76–79]. CPNA-1 has previously been shown to physically interact with DCR-1 [80], suggesting the potential regulation of *sago-2* by the RNAi machinery. However, a study has found that null mutations in *cpna-1* causes severe embryonic lethality and elongation defects [81], complicating efforts to study of its pleiotropic roles in RNAi and immunity.

While we found that SAGO-2 plays no role in transgene silencing in *pals-22(-)* mutants, it appears to target a subset of IPR genes (**Figure 5G**, **Figure S13**). Notably, as observed in *rrf-3* mutants, knocking out *sago-2* alone promotes IPR gene expression; however, eliminating *sago- 2* in the *pals-22(pig26)* background leads to a downregulation of IPR gene expression (**Figure 5G, H**). These results demonstrate that PALS-22 acts on distinct genomic targets via different WAGOs and suggest a potential gene-promoting role for SAGO-2 in the absence of PALS-22 (**Figure 5I**).

### Rapid evolution of the PALS-22/PALS-25 locus underlies natural variation in Mrt phenotype

Previous studies have shown that PALS-22 inhibits the IPR transcriptional program, which includes a suite of *pals* genes as well as the cullin-RING ubiquitin ligase components, promoting proteostasis capacities and resistance to infection with intracellular pathogens, such as *N. parisii* [32,37,71,82]. Building on these findings, we demonstrated that PALS-22 also inhibits the antiviral RNAi pathway, which functions to silence foreign genetic elements. Given the key roles of PALS- 22 in regulating innate immunity and RNAi – two highly evolutionary pliable systems – it is perhaps not surprising that the locus containing *pals-22* (and linked *pals-25*) is highly variable among *C. elegans* wild isolates [83]. This variability could stem from the host-pathogen evolutionary arm race, which shapes immune responses against different natural pathogens and facilitates niche adaption [83,84].

A recent study showed that many wild isolates growing at elevated temperatures (25°C) exhibit the Mrt phenotype, characterized by progressive loss of fertility, a phenomenon typically associated with dysregulation of small RNA inheritance and/or chromatin modifications. Interestingly, infection by microsporidia could rescue germline mortality in wild isolates, suggesting an interaction between the innate immunity and epigenetic inheritance processes [22]. Therefore, we asked whether PALS-22 might influence germline mortality and transgenerational effects. We found that natural variation in *pals-22/pals-25* locus in wild isolates correlates with the penetrance of the Mrt phenotype. Strains carrying alternate haplotypes resulting in loss of functions of *pals-22* and/or *pals-25* exhibit a more severe Mrt phenotype (a lower mean number of generations until they reach sterility at 25°C) compared to strains with the “wild-type” haplotype found the lab reference N2 strain (Mann Whitney test p = 0.0002; **Figure 6A**).

**Figure 6:**
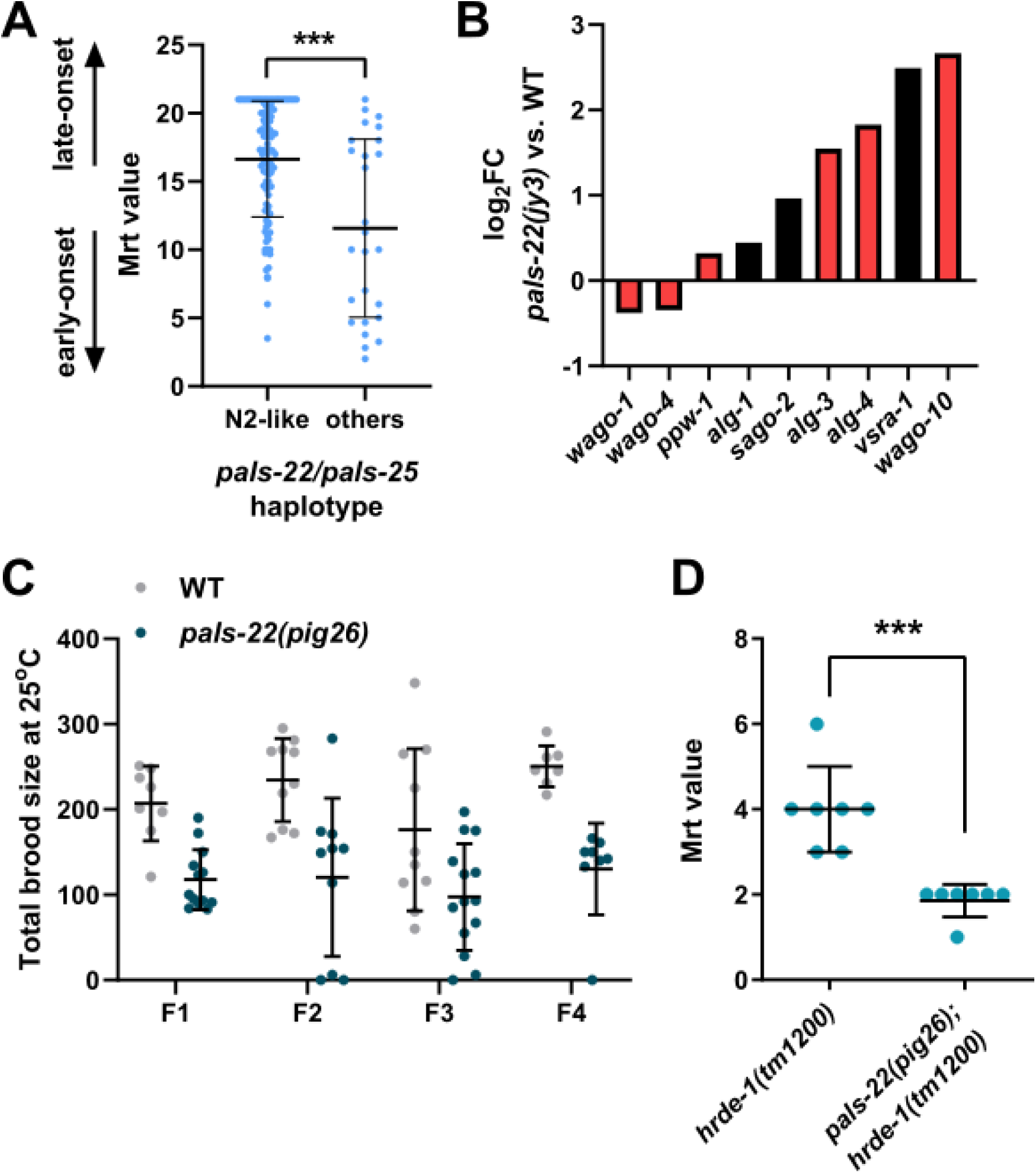
PALS-22 regulates germline mortality. (A) Mrt value (mean number of generations to sterility) in wild isolates carrying either N2-like or alternate haplotypes. Statistical significance was determined by non-parametric Mann-Whitney test. *** p ≤ 0.001. (B) Differential expression of *ago* genes in *pals-22(jy3)* mutants (L4/young adults) [39]. q < 0.01. Black and red bars indicate somatic and germline AGOs, respectively. (C) Total brood size of wild type and *pals-22(pig26)* mutants at 25°C. (D) Mrt value of *hrde-1(tm1200)* and *pals-22(pig26); hrde-1(tm1200)* mutants. Each dot represents a biological replicate. Statistical significance was determined by non-parametric Mann-Whitney test. *** p ≤ 0.001. Error represents mean +/− SD.

Similar to our findings on VSRA-1 and SAGO-2, we found that loss of *pals-22* causes dysregulation of multiple genes encoding for germline AGOs, including downregulation of WAGO- 1 and WAGO-4, which are known to play roles in epigenetic inheritance (**Figure 6B**). Although the loss of *pals-22* alone does not cause obvious germline mortality at 25°C (**Figure 6C**), eliminating *pals-22* severely enhances Mrt phenotype in *hrde-1* mutants, which encodes a germline WAGO that is important for small RNA inheritance (**Figure 6D**). These findings further strengthen the connections between the immune response and epigenetic inheritance.

In summary, our results demonstrate that PALS-22 is a rapidly evolving genetic node that controls multiple immunological programs against intracellular pathogens and genomic parasites. Natural variations in this key node likely influence how animals interact with their environments and affect their ability to transmit epigenetic information across generations. Hence, our results link host-pathogen co-evolution with transgenerational epigenetic inheritance - two processes that have major impacts on the evolutionary trajectory of the animals.

## Discussion

In this study, we reported four major findings: (1) PALS-22 represses the antiviral RNAi pathway, promoting the expression of repetitive transgenes and endogenous dsRNAs. (2) Genetic perturbations or infection by intracellular pathogens causes upregulation of VSRA-1 and SAGO- 1, which in turns regulate foreign genetic elements and immunity. (3) The endo-siRNA pathway normally silences IPR genes; however, in the absence of PALS-22, the endo-siRNA pathway upregulates immune gene expression. This finding suggests that PALS-22 functions as a molecular switch, dictating the functional states of the endo-siRNA pathway and highlighting a non-canonical gene-promoting role for small RNAs. (4) PALS-22 protects germline mortality, potentially by modulating the expression of germline AGOs, suggesting a role in epigenetic inheritance. Together, our findings suggest that PALS-22 modulates the regulatory output of RNAi to balance the trade-off between immunity against foreign elements and ‘autoimmunity,’ marked by aberrant silencing of self-genes and heritable germline defects. Natural variation in this critical genetic node may drive the evolution of alternative survival and reproductive strategies, facilitating niche adaptation.

Many previous studies on *C. elegans* AGOs have primarily focused on their roles in germline development and fertility. It was found that AGO localization to perinuclear liquid-like germ granules is critical for their functions in regulating germline gene expression homeostasis [85]. In the soma, subcellular localization of the nuclear RNAi factor NRDE-3 regulates the efficacy of RNAi [50]. Our findings suggest that the functions of AGOs may also be highly dependent on their spatiotemporal expression and stoichiometry. For example, eliminating *vsra-1* alone does not affect transgene silencing; however, when *pals-22* is eliminated, or upon viral infection, *vsra- 1* is upregulated ubiquitously. VSRA-1 is one of the few WAGOs that contain DEDH slicer domain and exhibit slicing activity *in vitro* [6,17]. It is possible that its slicing activity is uncovered only when its concentration reaches a critical threshold. In mammals, for example, the loading of miRNAs depends on the expression levels of the different AGOs [86]. In *C. elegans*, a recent study showed that the expression levels of ALG-1 and ALG-2, which function in the miRNA pathway, strongly affect miRNA processing and animals’ viability. Notably, overexpression of slicing dead ALG-1/2 in *alg-1/2(-)* mutant backgrounds does not phenocopy animals with mutations in the catalytic tetrad but retains endogenous expression levels [87,88], suggesting that AGO functions are sensitive to molecular stoichiometry.

Most interestingly, our results indicate a potential cryptic gene-promoting role for SAGO- 2, acting in the RRF-3-dependent endo-siRNA pathway, when *pals-22* is eliminated. The molecular mechanism underlying this gene-promoting activity of SAGO-2 is currently unclear. One possibility is that the endo-siRNA pathway may enhance the expression of IPR genes indirectly by silencing an IPR suppressor. However, our observation of increased small RNAs targeting IPR genes in *pals-22(-)* mutants hints at a more direct mechanism. An alternative explanation might be that, in wild-type background, low level of SAGO-2 silences IPR genes. Note that SAGO-2 lacks the DEDH catalytic tetrad in its PIWI domain and cannot directly initiate degradation of its mRNA targets [6]; thus, involvement of an additional endonuclease is likely. In the absence *pals- 22*, *sago-2* is upregulated, the elevated level of SAGO-2 may alter its functions: instead of downregulating IPR transcripts, it may act as a “sink”, binding to and protecting them from endonucleases, thereby promoting the expression of IPR genes. Such a mechanism could exponentially expand the complexity of RNAi regulatory outputs, particularly in the face of environmental stress or genetic perturbation, warranting many follow up studies.

Together with previous studies, our results indicate that PALS-22 modulates multiple immunological programs – the transcriptional IPR and the antiviral RNAi. Innate immunity and RNAi tend to be rapidly evolving processes as they are locked in a tight arm race against pathogens and genomic parasites. PALS-22, as a common regulator in the two systems, may serve as a particularly plastic node for tuning the animals’ defense in response to environmental stimuli. Indeed, the *pals* gene family is rapidly evolving. The *pals-22* (and its neighboring *pals-25*) contains extensive polymorphisms among *C. elegans* wild isolates [83]; PALS-22 and PALS-25 have no obvious orthologs in other closely related *Caenorhabditis* species [39,43]. This natural variation likely affects how the animals interact with pathogens in the wild. Interestingly, a recent study has shown that associating *C. elegans* wild isolates with their natural microbes and microsporidia could rescue heat-induced mortal germline phenotype caused by transgenerational inheritance of aberrant epigenetic signals [22]. We found that the loss of *pals-22* enhances Mrt phenotype in wild isolates and small RNA inheritance mutants, supporting a role of PALS-22 in coordinating the immune response and the epigenetic inheritance machinery consisting of a suite of germline AGOs, which appear to be misregulated in animals lacking *pals-22.* To our best knowledge, this is one of the first reports to have mapped a genetic circuit that dynamically controls AGO expression and tissue specificity in animals.

We and others have shown that PALS-22 controls many life history traits, including pathogen resistance [39,71], reproduction, and lifespan [43], which have direct impact on the animals’ competitive fitness. PALS-22 and PALS-25 function as ‘antagonistic paralogs’: PALS-22 acts as a repressor, while PALS-25 serves as an activator of IPR, together balancing pathogen resistance, growth, and aging [39]. Other *pals* genes (*pals-17*, *pals-20*, and *pals-26*) have also been shown to regulate similar tradeoffs between immunity and development [89]. Curiously, a recent study showed that knocking out cGAS in mice causes depression of LINE1 transposons, induction of inflammation, and accelerated aging [90]. This suggests a conserved link between immunity and genomic surveillance mechanisms and reveals a surprising parallel between the highly divergent innate immune systems of *C. elegans* and mammals. Indeed, the gene regulatory architecture of IPR resembles that of type-I interferon response [32]. Although the functions of many *pals* genes have not been studied, it is tempting to speculate that copy number of *pals* genes may underlie rapid diversification of the gene regulatory network governing RNAi and innate immunity, and contribute to life history evolution in nematodes.

## Methods

### *C. elegans* cultivation

*C. elegans* strains were propagated at 20°C in standard Nematode Growth Media (NGM) seeded with *E. coli* OP50 [91], unless stated otherwise. Worm strains and primers used in this study were listed in Tables S2 and S3, respectively.

### RNAi

RNAi feeding clones were obtained from the Ahringer or the Vidal libraries [92,93]. The bacteria were inoculated overnight at 37°C in LB containing 50 μg/ml carbenicillin. The next day, 1 mM of IPTG was added to the bacterial culture and 50-100 μL was seeded onto 60 mm NGM agar plates containing 1 mM IPTG and 25 μg/mL carbenicillin. After the seeded plates were dried for about 24 hours, 5-10 L4 animals were placed on the RNAi plates. The progeny were then collected for analyses.

### Imaging and fluorescence quantification

The animals were immobilized with 5 mM levamisole and mounted on 2% agarose pads. Fluorescence Images were taken using Olympus IX83 motorized inverted wide-field microscope or Olympus BX63 motorized upright wide-field microscope. Transgene expression was analyzed using Fiji/ImageJ software and genotypes were blinded from the investigator using the software DoubleBlind (https://github.com/GuyTeichman/DoubleBlind). Corrected total worm fluorescence = integrated density – (area of selected cell X mean fluorescence of background readings). Overlapping worms and worms on edges of the image were discarded from analysis.

### RNA sequencing

Gravid adults were treated with alkaline hypochlorite solution and the embryos were thoroughly washed and incubated in M9 overnight to obtain a synchronized L1 population. Total RNA was isolated from L1 animals using standard phenol-chloroform method using TRIzol or Qiagen RNeasy Kits per manufacture instruction.

Libraries for RNA-seq were prepared using NEBNext® Ultra II Directional RNA Library Prep Kit for Illumina® coupled with NEBNext® Poly(A) mRNA Magnetic Isolation Module. The sample quality was accessed on Agilent 2200/4150 BioAnalyzer instrument and High Sensitivity RNA ScreenTapes. The resulting cDNA libraries were pooled and paired-end sequencing was performed on the Nextseq 550/2000 platform.

For small RNA-seq, the total RNA was first treated with RNA 5′ Polyphosphatase (epicentre) followed by library preparation using NEBNext® Small RNA Library Prep Set for Illumina®. Size selection on E-gel (Invitrogen, Life Technologies) was performed to enrich for 140- 160 nt long cDNA. The sample quality was then accessed on Agilent 2200/4150 BioAnalyzer instrument and High Sensitivity RNA ScreenTapes. The sampled were pooled and sequenced on the Nextseq 550/2000 platform.

### Bioinformatics

All bioinformatic analyses were performed using RNAlysis [94] (https://github.com/GuyTeichman/RNAlysis). For small RNA sequencing datasets, we filtered out reads <15 and >30 nucleotides after adapter trimming using CutAdapt [95]. We aligned small RNA reads to PRJNA13758 ce11 genome assembly using ShortStack [96] and counted aligned sense and anti-sense small RNA reads using FeatureCounts (**Supplementary File 1**) [97]. For mRNA sequencing datasets, we pseudo-aligned reads using Kallisto (**Supplementary File 2**) [98]. We performed differential expression analysis using DESeq2 [99]. For gene set enrichment analysis, log_2_ (fold enrichment) scores were calculated and the FDR for enrichment was calculated using 10,000 random gene sets identical in size to the tested group.

### qPCR

mRNA was isolated from L1 animals using Qiagen RNeasy Kits as described above. cDNA was generated using High-Capacity cDNA Reverse Transcription Kit (Applied Biosystems) and qPCR was performed using 2x qPCRBIO SyGreen Blue Mix Lo-ROX. The 2–ΔΔCt method was used to calculate the relative fold-change of the samples. qCPR primers used are provided in Table S3.

### CRISPR/Cas9 gene editing

To generate *pals-22(pig26)* allele, we used two guide RNAs (TGGCCTGCAAGACGAGGTAC and GGTTTGGGAAGAACTTAAAA) and a repair template (CAGGTTGCATAGAGAAGAAGATGAACTCGCCGGTAAAAAGGTGTTGGAAGCGGCTGAAATAGTTGATGTG) ordered as a DNA gBlock from Integrated DNA Technologies (IDT). CRISPR/Cas9 edits were performed as previously described [100]. The resulting mutant carrying 253 bp deletion in *pals-22* was backcrossed to N2 twice to eliminate background mutations.

### Mortal germline assay

Mortal germline assay was performed as previously described [22,101]. Briefly, four L4 animals were transferred from 20°C to 25°C (labeled F1). At every generation, four L4 animals were transferred to fresh seeded NGM plate until the population extinct. The number of generations to sterility was recorded.

### Movement tracking

20-30 L4 worms were placed on NGM plates seeded with OP50. The worms were recorded on the following three days. Each day, they were recorded for a duration of 5 mins, capturing two frames per second using Teledyne Dalsa M2420 monochrome camera. Worm motility was subsequently analyzed using the Fiji/ImageJ wrMTrck plugin (http://www.phage.dk/plugins/wrmtrck.html).

### Brood size assay

Transfer individual L4 animal onto NGM plates seeded with OP50. The worms were transferred to fresh plates daily and the number of progeny laid each day was scored for 4-5 days, until the worms are sperm depleted.

### *N. parisii* and Orsay virus infection assays

*N. parisii* spores were isolated as previously described [84]. 1200 synchronized L1 animals were mixed with 5.5 × 10^5^ *N. parisii* spores and 25 µL of 10X concentrated OP50-1 bacteria, and then suspended to a total volume of 300 µL in M9 buffer. This mixture was top-plated onto unseeded 6 cm NGM plates in duplicate per condition, allowed to dry for 30 minutes at RT, and then incubated at 25 °C for a 3-hour infection. Worms were washed off plates and fixed in 4% PFA prior to incubation with a FISH probe specific to *N. parisii* ribosomal RNA conjugated to a Cal Fluor 610 fluorophore (Biosearch Technologies). Pathogen load was determined by quantifying the number of sporoplasms present per animal across 100 worms per condition. Quantification was performed using an AxioImager under a 40X objective lens.

Infectious Orsay virus filtrate (OVF) was prepared as previously described [102]. 2000 synchronized L1 animals were plated onto 10 cm NGM plates seeded with 500 µL of 10X OP50- 1 and allowed to grow to L4 stage (44 hours at 20 °C), after which an infection mix comprised of 150 µL of 10x OP50-1, 30 µL of OVF, and 720 µL of M9 (900 µL total per plate) was top-plated. Once dry, plates were incubated at 20 °C for 24 hours prior to fixation in 4% PFA and incubation with two FISH probes specific to both Orsay virus RNA1 and RNA2, each conjugated to a Cal Fluor 610 fluorophore (Biosearch Technologies). Images of the FISH probe signal across approximately 15 worms per condition were taken using an AxioImager under a 10X objective, and the signal was then quantified using ImageJ, correcting for background fluorescence.

### Statistics and figure preparation

Statistics were performed using R software 4.2.3 (https://www.r-project.org/) and GraphPad Prism 8. The specific statistical tests are reported in the figure legends. Plots were generated using GraphPad Prism 8. Figures were assembled in Inkscape v0.92.4 (https://inkscape.org/) or BioRender (http://www.BioRender.com/)

## Data Availability

All NGS data are available through GEO, accession number [TBA]. The following previously published datasets were used in this study: GSE143595, GSE208702, SRP100798, and GSE118400.

## Acknowledgment

We thank Michalis Barkoulas (Imperial College London) for helpful comments on the manuscript. O.R. is grateful to funding from the Eric and Wendy Schmidt Fund for Strategic Innovation (Polymath Award #0140001000) and the generous support from the Morris Kahn foundation. G.T. is grateful to the support from Milner Foundation. M.M.L.K. is supported by the Azrieli Foundation and C.K.E. is supported by EMBO Postdoctoral Fellowship (#ALTF 6-2022). This work is supported by ERC grant #335624 to O.R. and NSF 2301657 to E.R.T. We are grateful to WormBase for providing valuable data and resources. Some strains were provided by the CGC, which is funded by NIH Office of Research Infrastructure Programs (P40 OD010440).

## Supplementary data

**Figure S1:**
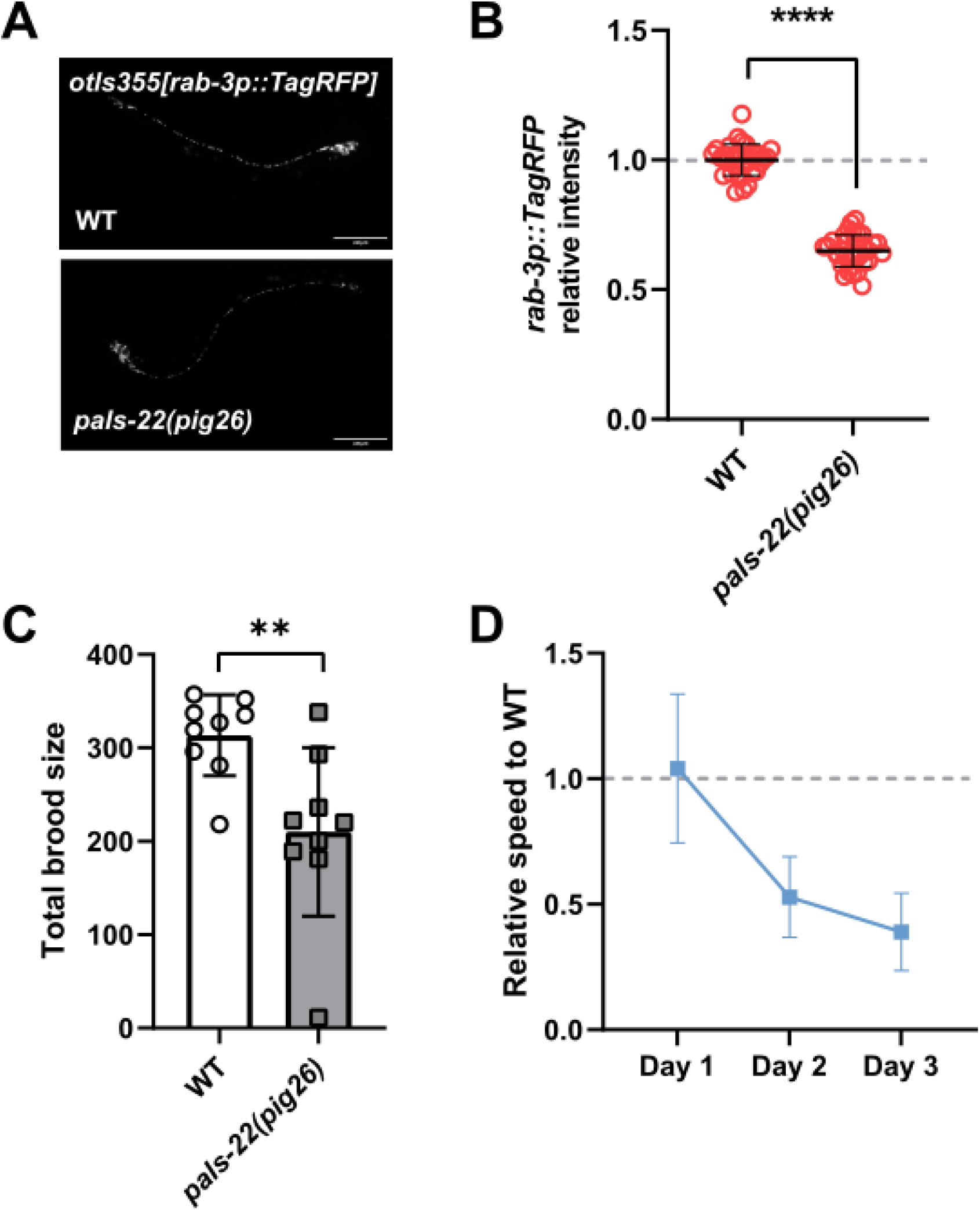
(A, B) Expression of *otIs355[rab-3p::TagRFP]* in WT and *pals-22(pig26)* mutants. Scale bar = 100 µm (C) Brood size of wild type and *pals-22(pig26)* mutants at 20°C. (D) Locomotion speed of *pals-22(pig26)* mutants normalized to aged matched wild type animals. Statistical significance was determined by two-tailed unpaired t-tests. ** ≤ 0.01; **** p ≤ 0.0001.

**Figure S2:**
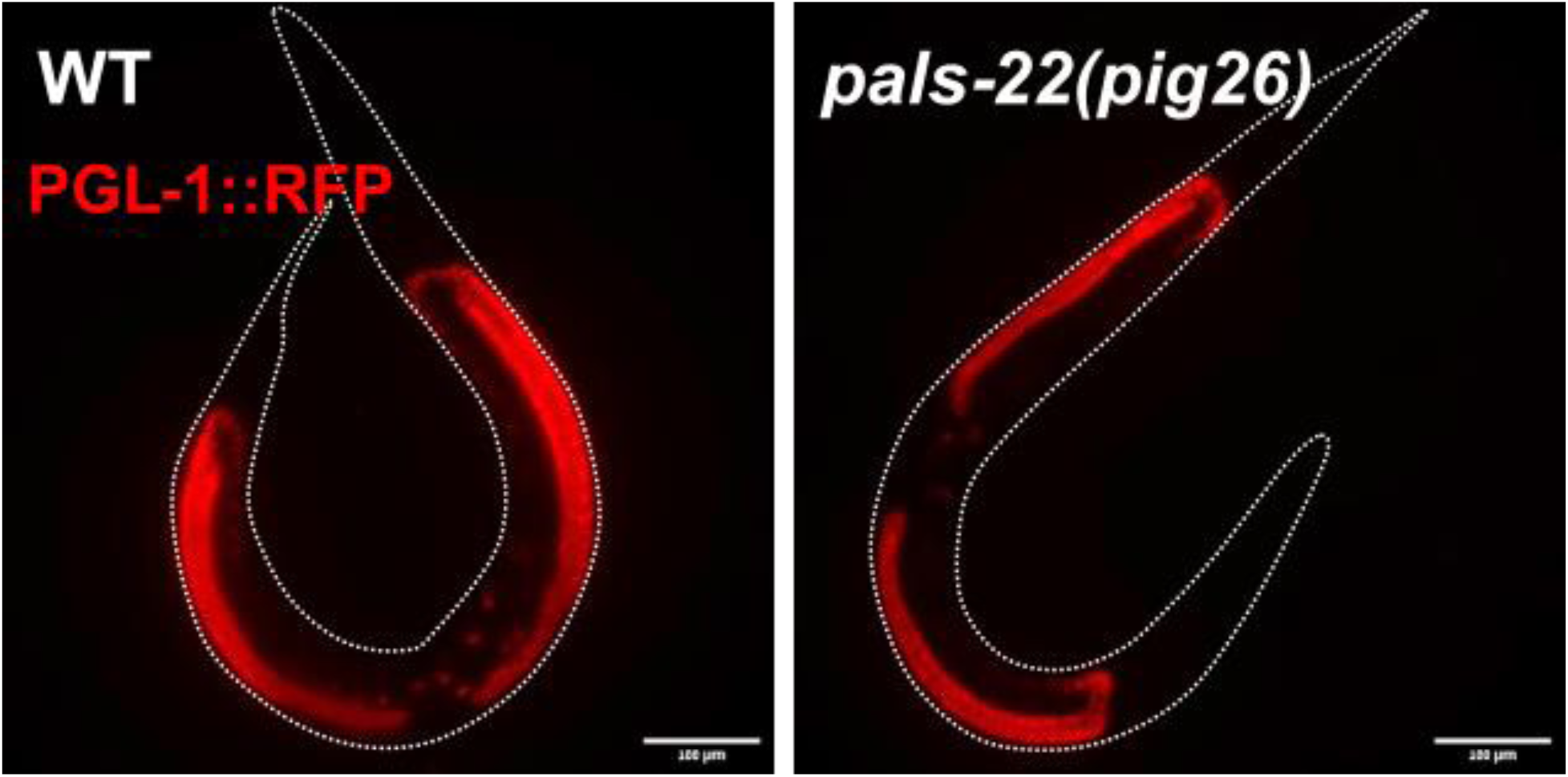
The expression of *pgl-1::TagRFP* in wild type and *pals-22(pig26)* day 1 adults. Scale bar = 100 µm.

**Figure S3:**
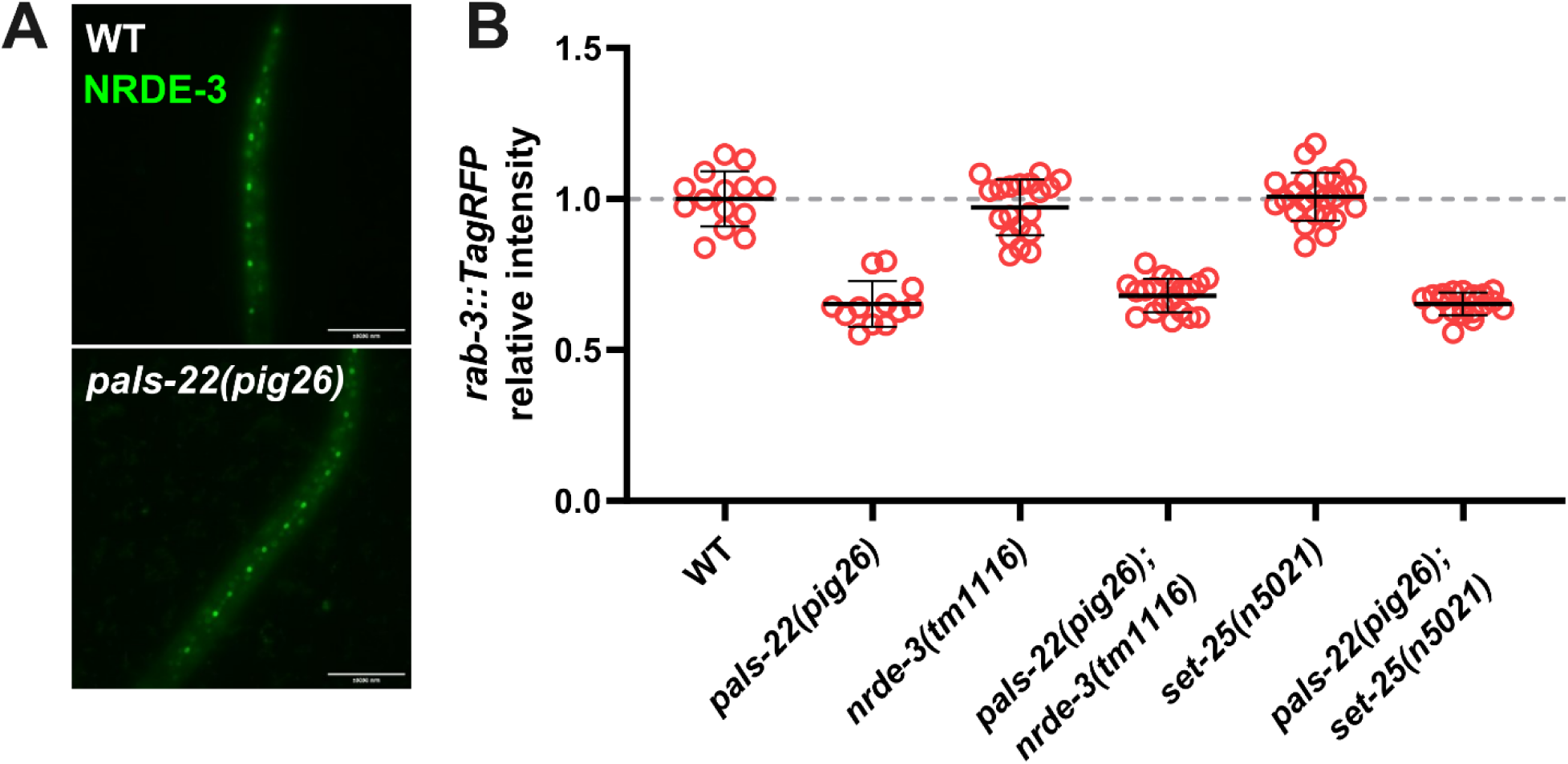
(A) The expression of endogenous NRDE-3::GFP reporter in wild-type and *pals- 22(pig26)* L2 animals. Scale bar = 10 µm. (B) The expression repetitive *rab-3::TagRFP* transgene in *pals-22(pig26)* and nuclear RNAi mutants. Each data point represents an animal scored.

**Figure S4:**
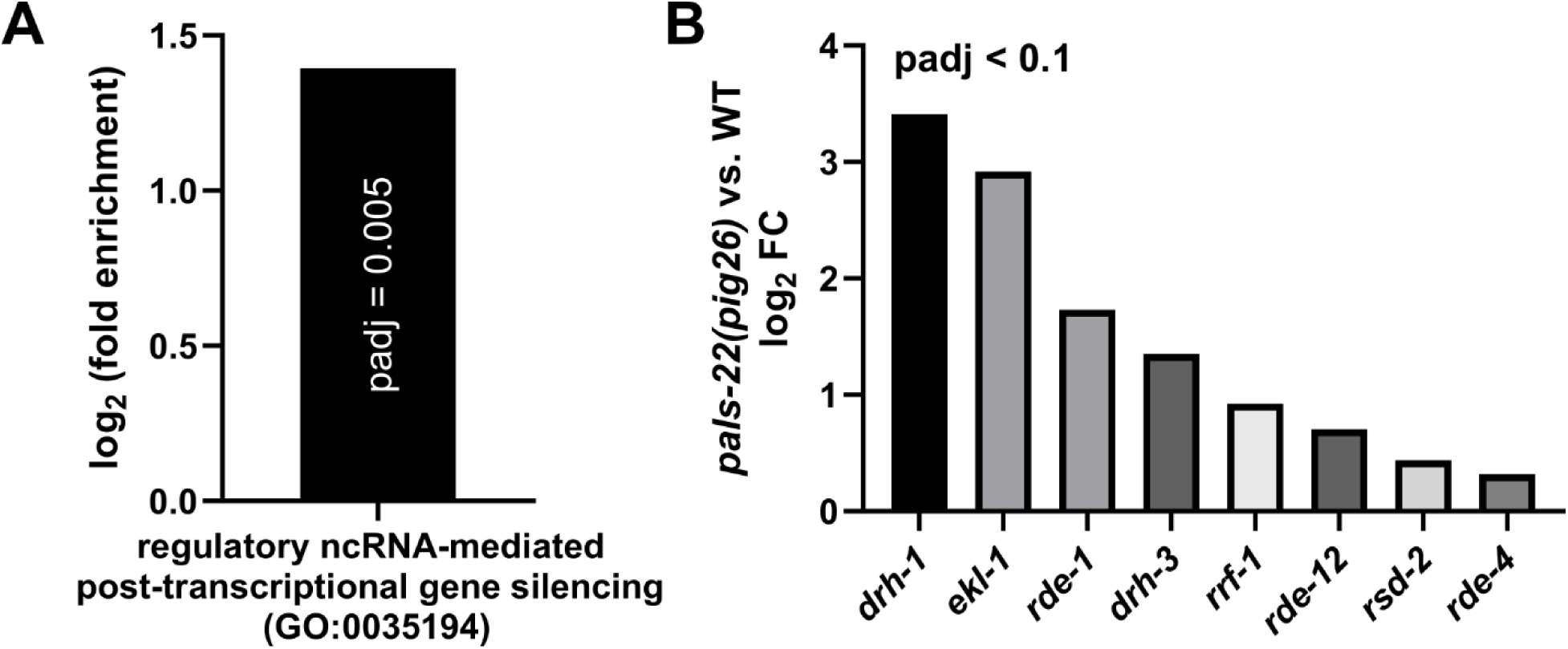
(A) GO term enrichment of genes upregulated in *pals-22(pig26)* vs. wild type. (B) RNAi genes upregulated in *pals-22(pig26)* vs. wild type.

**Figure S5:**
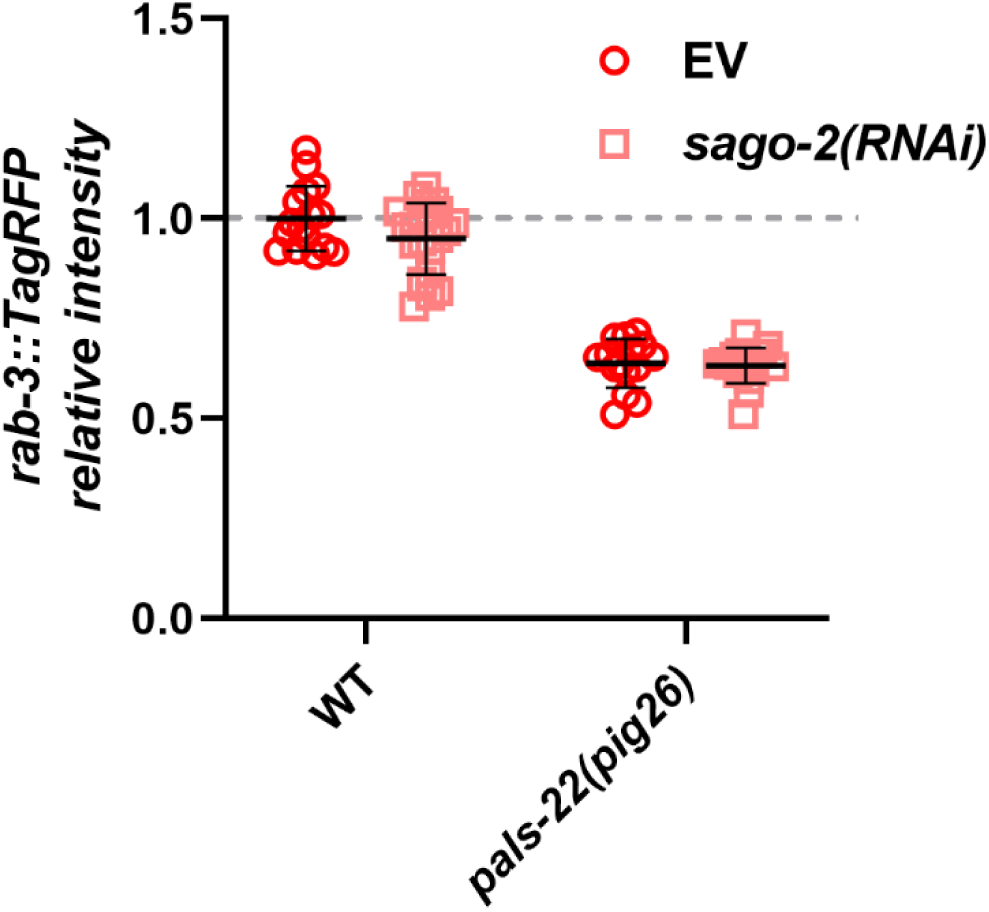
(A) The effects of *sago-2(RNAi)* on transgene expression in wild type and *pals- 22(pig26)*. Each data point represents an animal scored.

**Figure S6:**
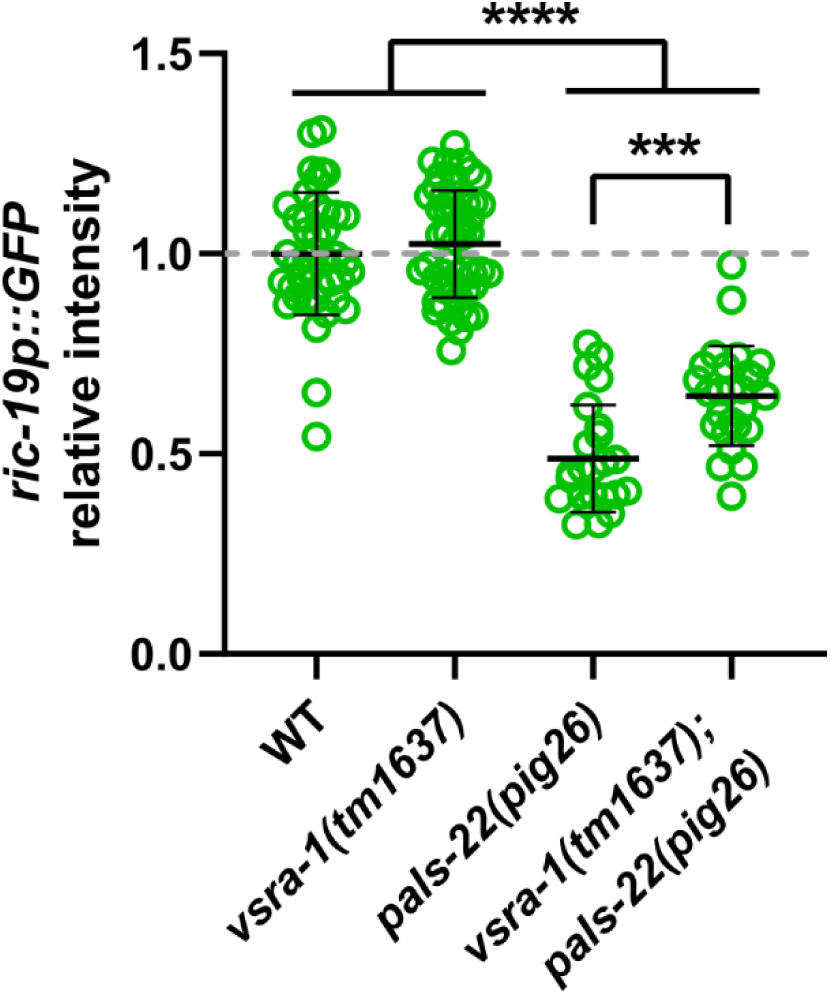
The interaction between *pals-22* and *vsra-1* in silencing *otIs381 [ric- 19p(prom6)::2xNLS::GFP + elt-2::DsRed].* Statistical significance was determined by parametric one-way ANOVA followed by pairwise two-tailed unpaired t-tests with Benjamini–Hochberg correction. *** q ≤ 0.001; **** q ≤ 0.0001. Error represents mean +/− SD.

**Figure S7:**
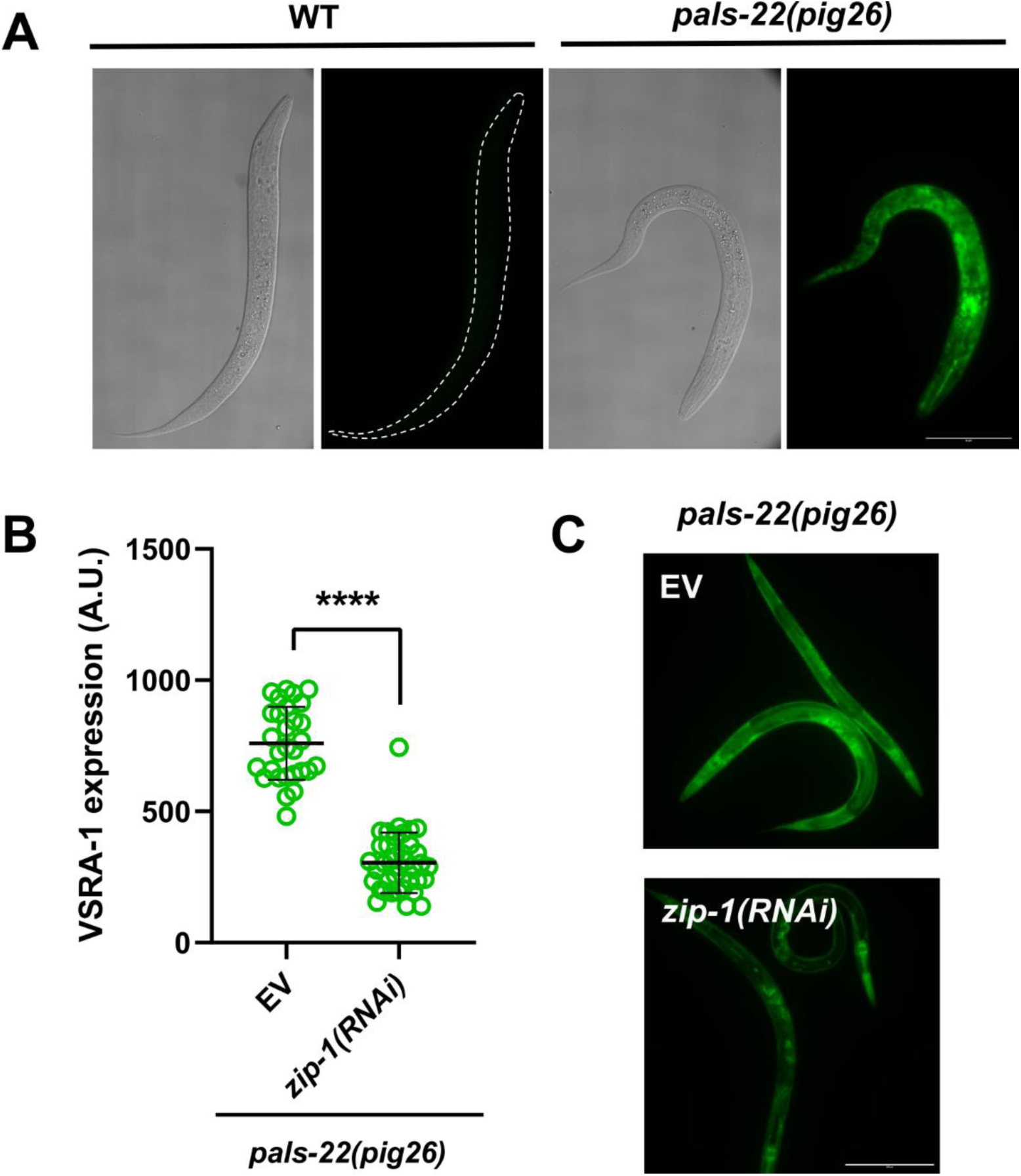
(A) The expression of endogenous VSRA-1 in wild-type and *pals-22(pig26)* L1 animals. Scale bar = 50 µm. (B, C) The expression of VSRA-1 in *pals-22(pig26)* animals treated with empty vector or *zip-1* RNAi. Statistical significance was determined by two-tailed unpaired t-test. **** q ≤ 0.0001. Error represents mean +/− SD. Scale bar = 200 µm.

**Figure S8:**
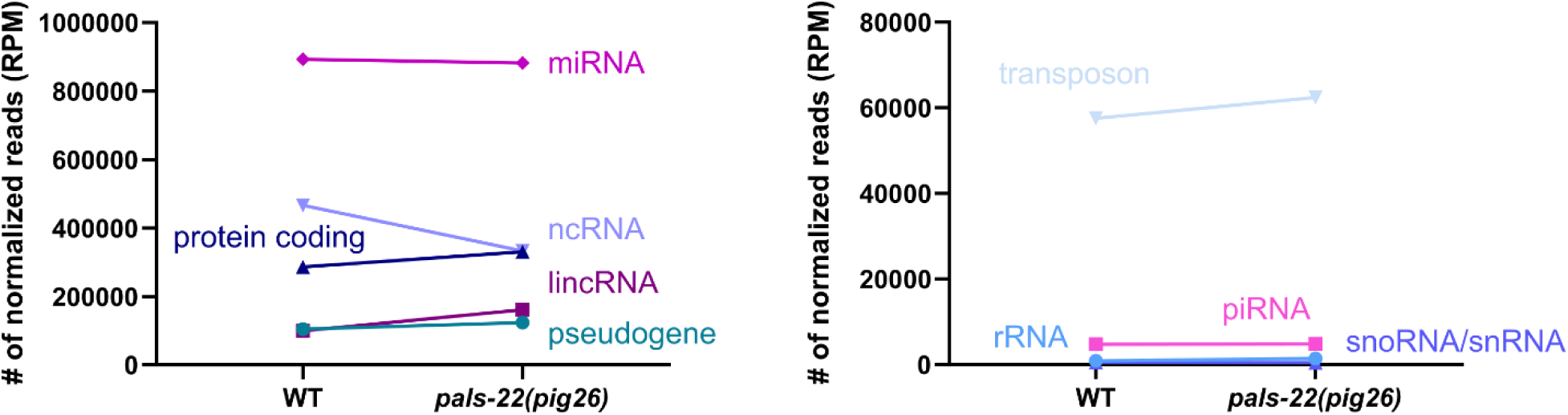
Biotypes of small RNAs isolated from wild-type and *pals-22(pig26)* mutants. RPM: read per million.

**Figure S9:**
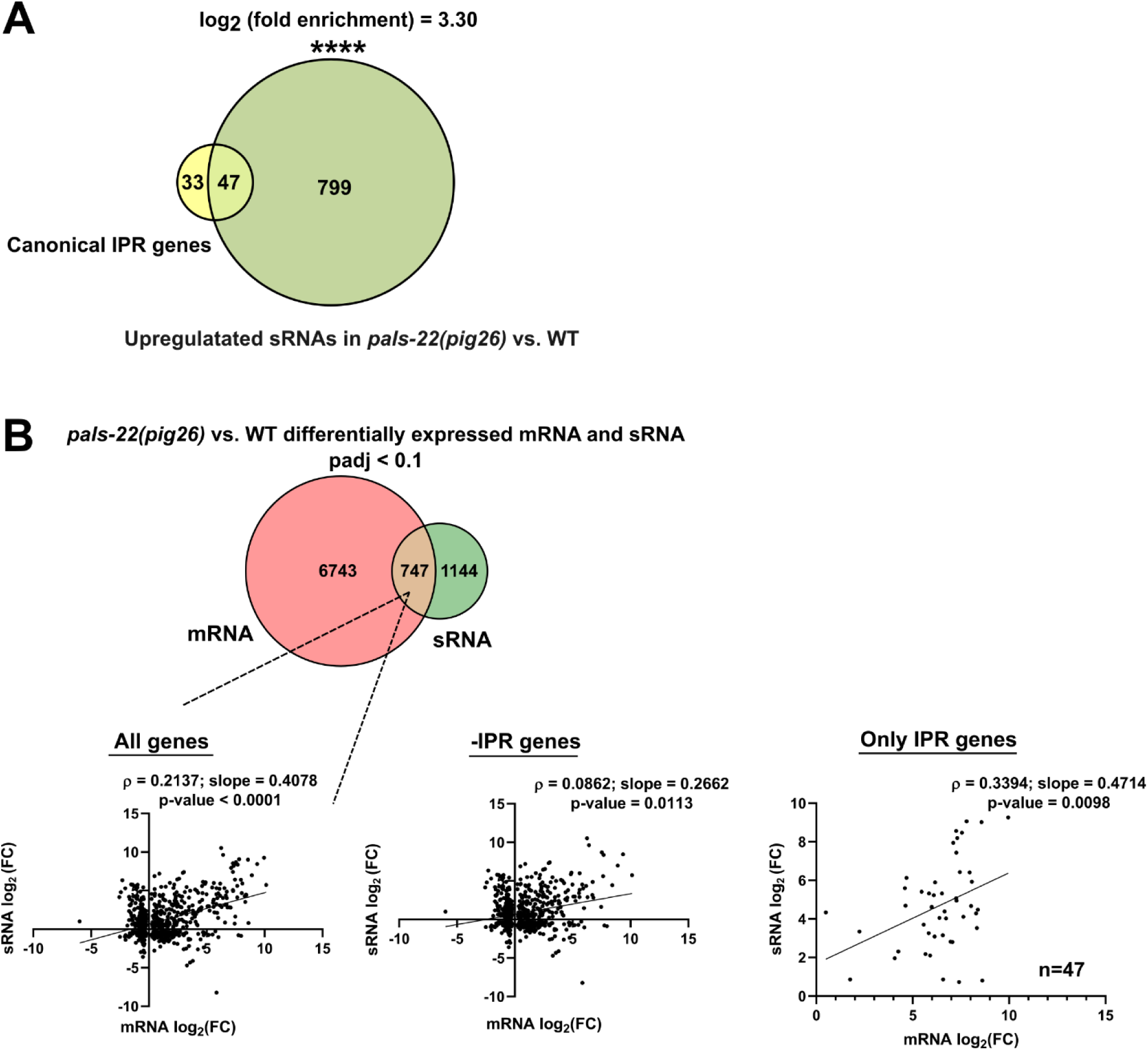
(A) Venn diagram showing the overlap between 80 canonical IPR genes and small RNA upregulated in *pals-22(pig26)* vs. wild type. (B) Spearman correlation between differentially expressed mRNA and small RNA in *pals-22(pig26)* vs. wild type. The lines are fitted with linear regression model.

**Figure S10:**
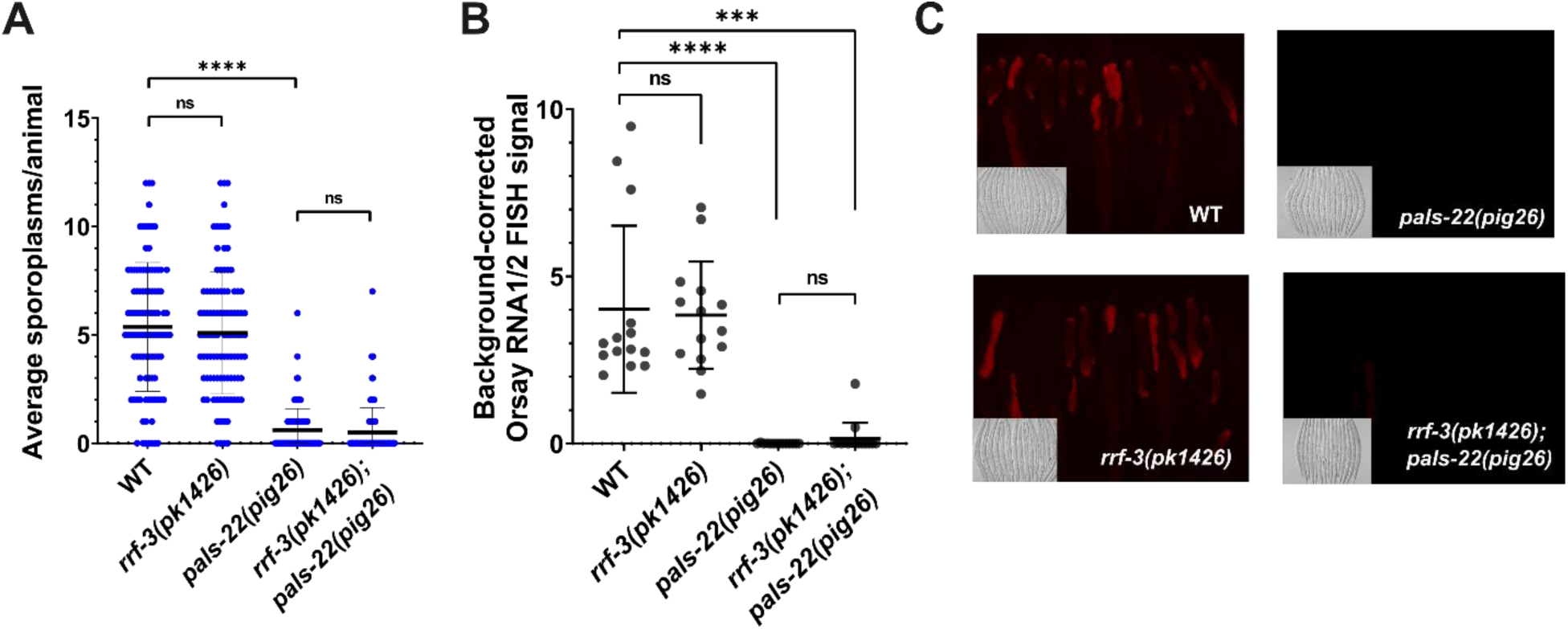
(A) Sensitivity of *rrf-3(pk1426)*, *pals-22(pig26)*, and *rrf-3(pk1426); pals-22(pig26)* mutants to *N. parisii* infection. (B) Sensitivity of *rrf-3(pk1426)*, *pals-22(pig26)*, and *rrf-3(pk1426); pals-22(pig26)* mutants to Orsay virus infection. Statistical significance was determined by non-parametric Kruskal–Wallis test followed by pairwise Wilcoxon Rank Sum tests with Benjamini Hochberg correction. ns, not significant (q > 0.05); *** q ≤ 0.001; **** q ≤ 0.0001. Error represents mean +/− SD. (C) Representative micrographs of animals from (B).

**Figure S11:**
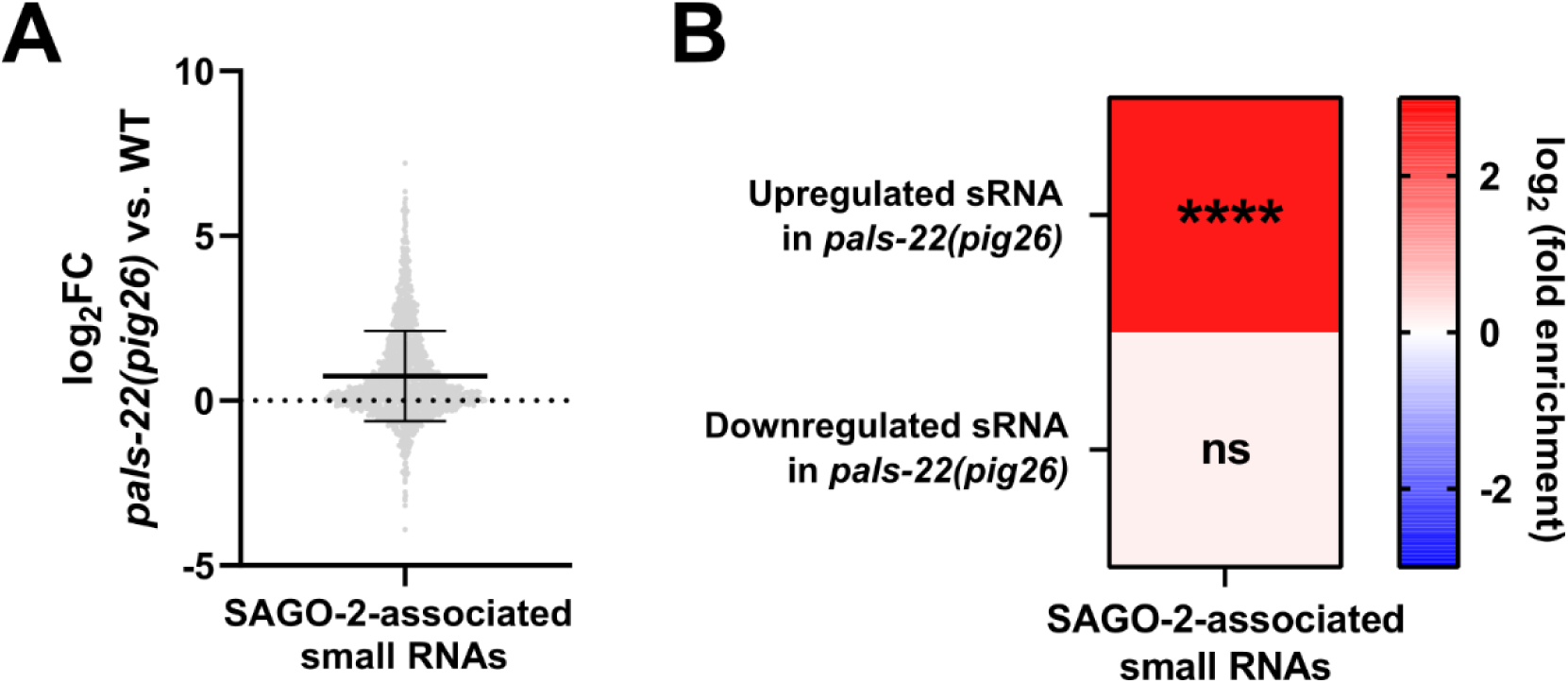
(A) The expression of SAGO-2-associated small RNAs in *pals-22(pig26)* vs. wild type. (B) Enrichment analysis of SAGO-2-associated small RNAs in small RNAs differentially expressed in *pals-22(pig26)* vs. wild type.

**Figure S12:**
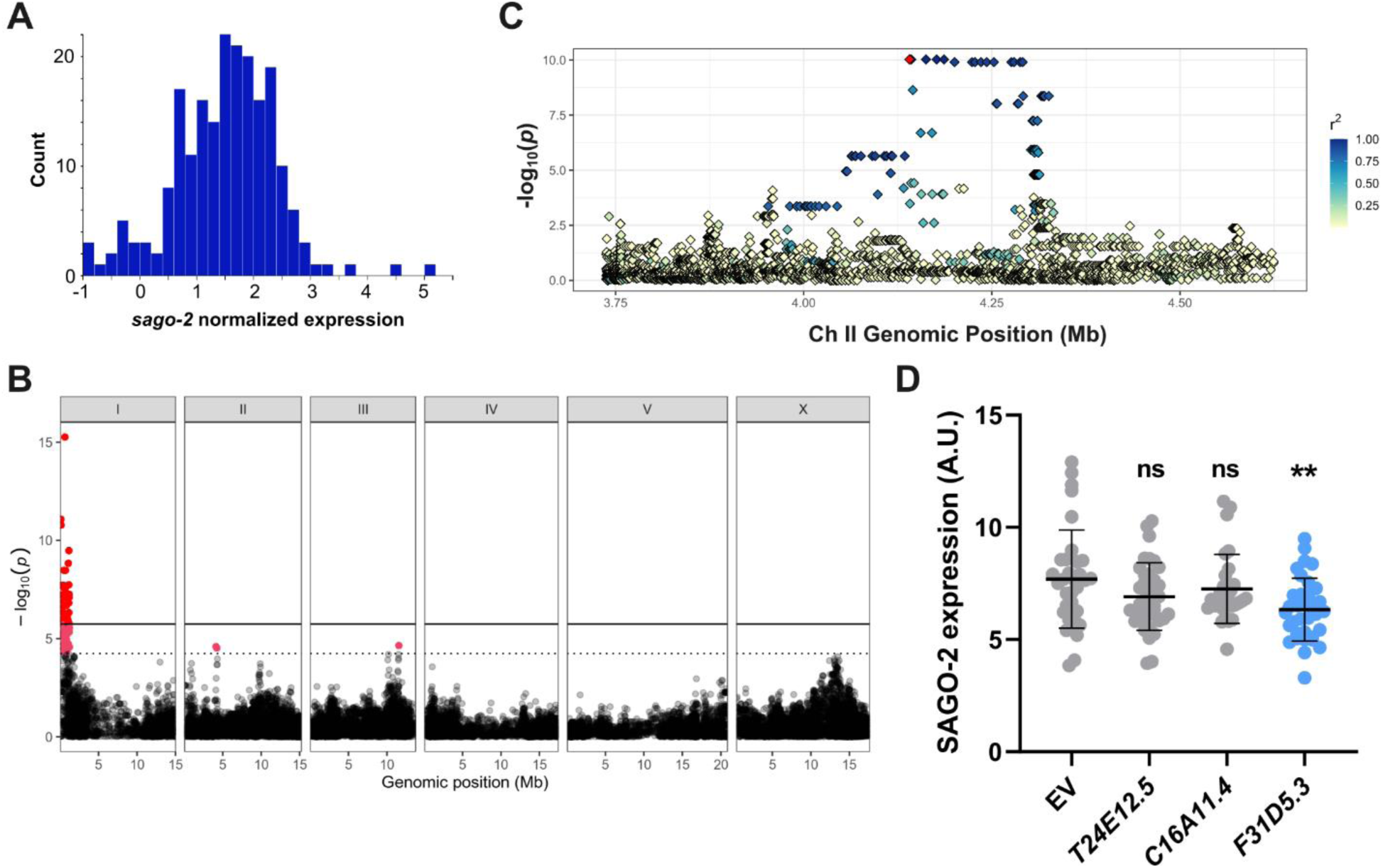
(A) Natural variation of *sago-2* expression in 207 *C. elegans* wild isolates. (B) GWAS using LOCO kinship matrix. The horizontal solid line corresponds to stricter Bonferroni (BF) threshold, while the horizontal dash line corresponds to more permissive EIGEN threshold. Red dots represent statistically significant SNPs. (C) Fine mapping of Chr 2 candidate region (II:3737765-4624311). Each variant is represented by a diamond colored by the linkage to the peak marker (colored in red). GWAS was performed on CaeNDR (https://caendr.org/). (D) Targeted RNAi screen focusing on candidate genes in Chr 2. Statistical significance was determined by non-parametric Kruskal–Wallis test followed by pairwise Wilcoxon Rank Sum tests with Benjamini–Hochberg correction. ns, not significant (q > 0.05); ** ≤ 0.01, compared to EV.

**Figure S13:**
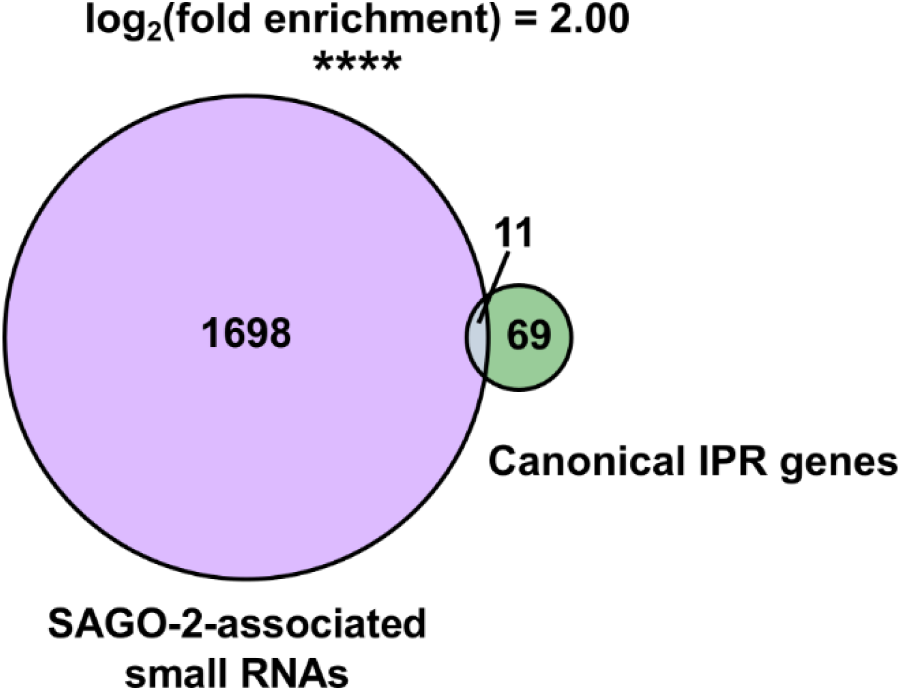
Venn diagram showing the overlap between 80 canonical IPR genes and SAGO-2- associated small RNAs.

**Table S1:** Expression of *vsra-1* based on single-cell sequencing of young adult worm (WormSeq [103]). Only tissues with detectable expression are shown below. TPM = transcript per million.

| Tissue | Scaled TPM |
| --- | --- |
| Unassigned sheath cells | 90.95 |
| AVL | 57.83 |
| Glia_3 | 54.23 |
| sh3_sh4 (gonadal sheath proximal) | 52.94 |
| Oocytes | 44.99 |
| Amphid sheath | 37.31 |
| AWB | 36.56 |
| Spermathecal-Uterine junction | 36.11 |
| Coelomocytes | 30.49 |
| RIR | 23.35 |
| Phasmid sheath | 22.33 |
| Dorsal uterine cell | 19.45 |
| AVM | 18.95 |
| RMD_LR | 8.90 |
| Intestinal-rectal valve | 4.36 |
| g2 | 2.30 |
| Spermathecal bag distal | 1.12 |

**Table S2:**
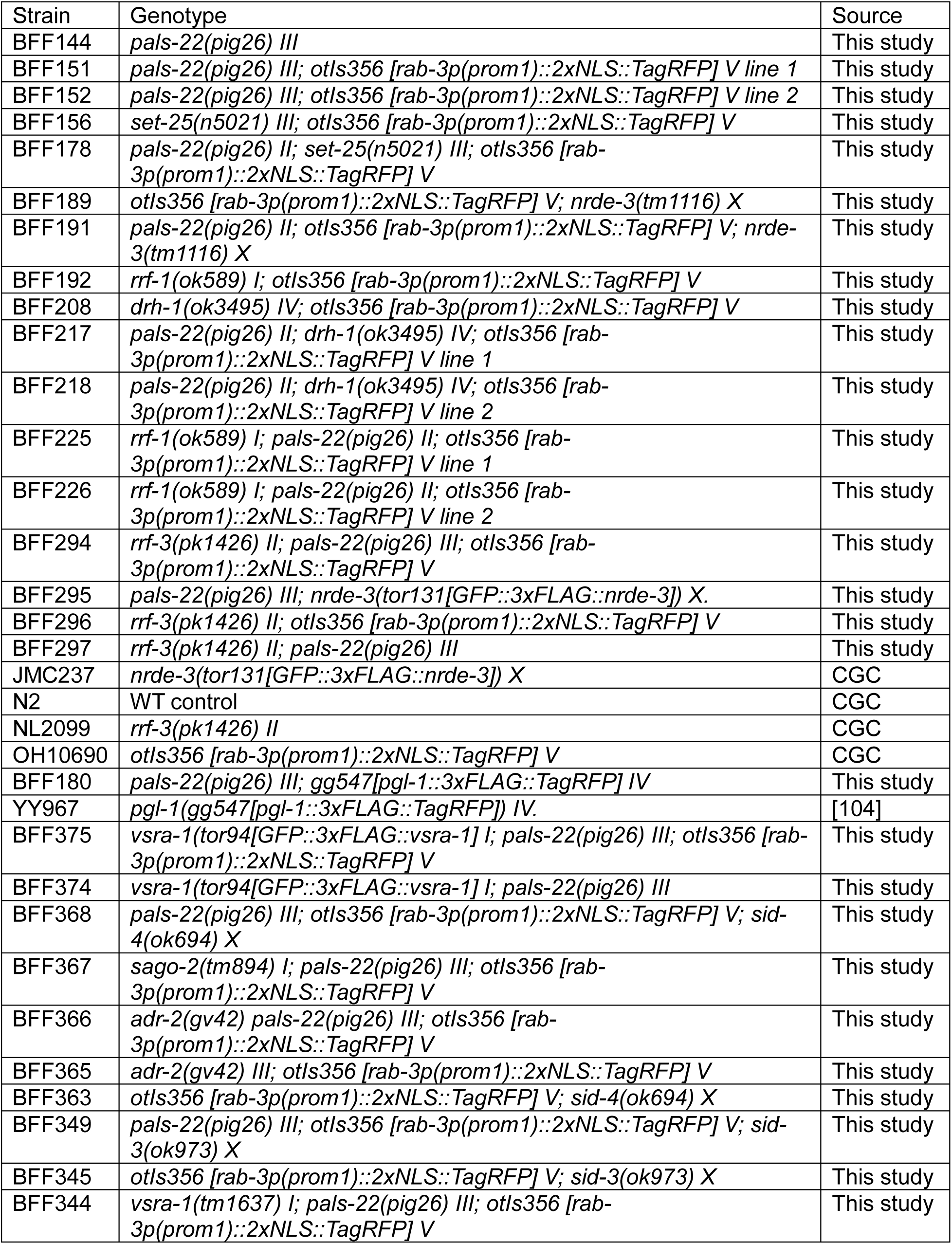
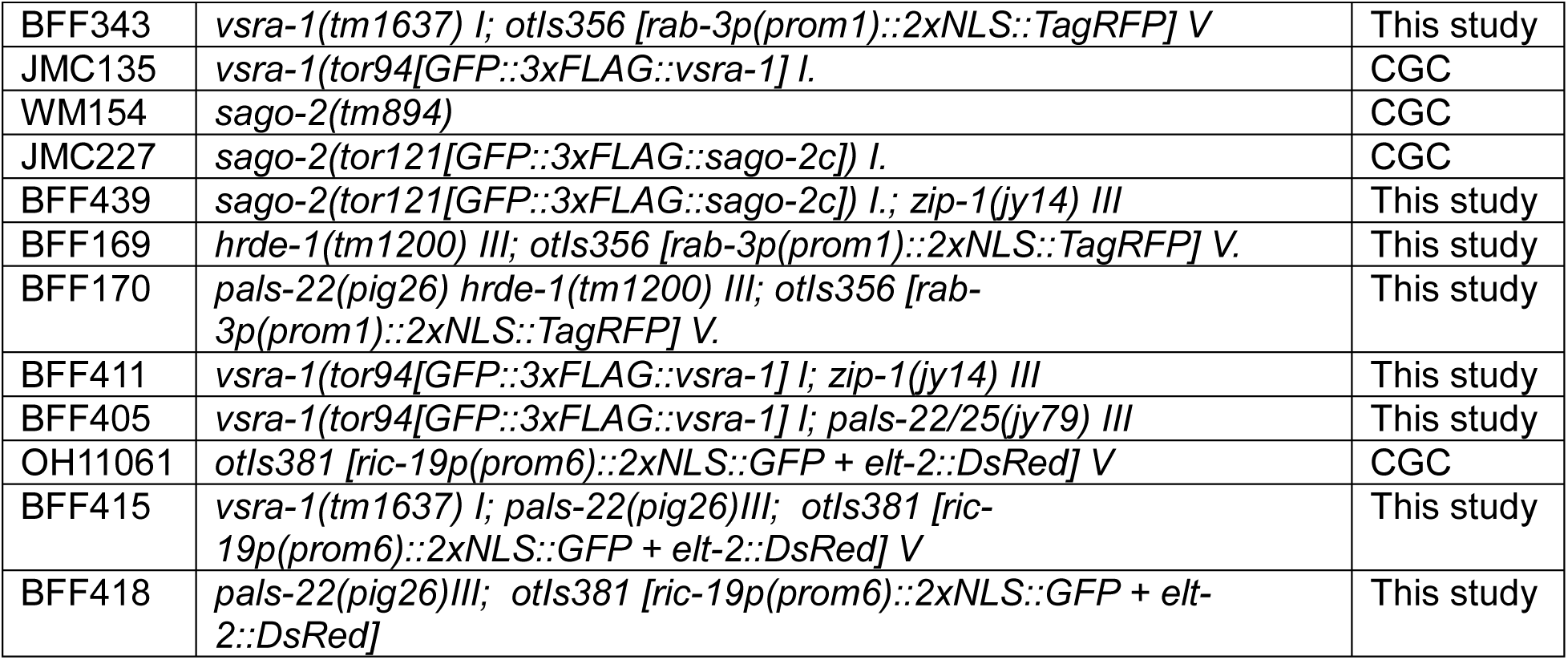
Worm strains used in this study.

**Table S3:**
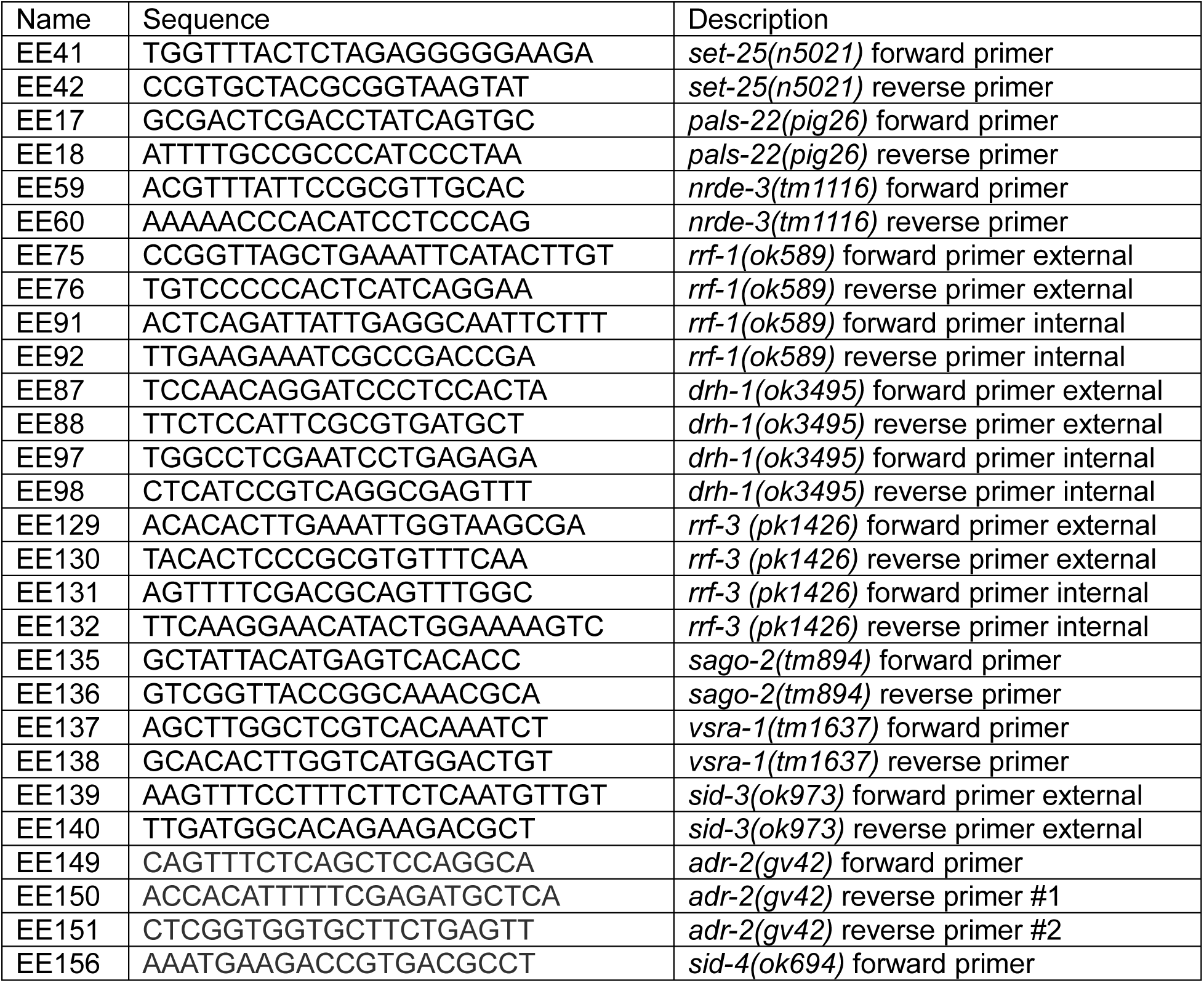
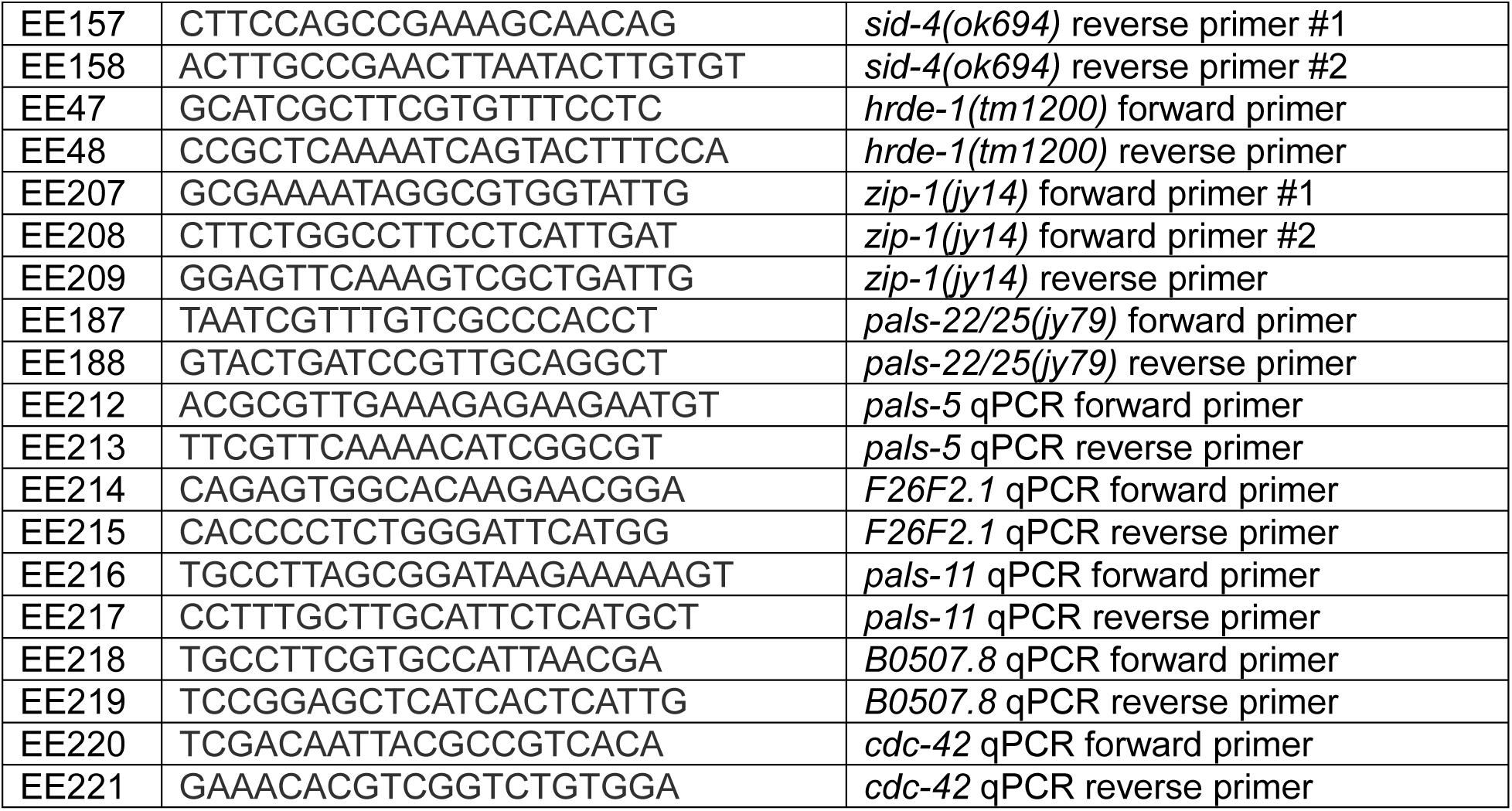
Genotyping primers used in this study.

**Supplementary File 1 (xlsx):** FeatureCounts and DESeq2 outputs of antisense small RNA in wild type and *pals-22(pig26)*.

**Supplementary File 2 (xlsx):** Kallisto and DESeq2 outputs of mRNA in wild type, *pals- 22(pig26), rrf-3(pk1426)*, and *rrf-3(pk1426); pals-22(pig26)*.

## References

1. Seroussi U, Lugowski A, Wadi L, Lao RX, Willis AR, Zhao W, et al. A comprehensive survey of C. elegans argonaute proteins reveals organism-wide gene regulatory networks and functions. James DE, Pasquenelli A, Kennedy S, editors. eLife. 2023;12: e83853. doi:10.7554/eLife.83853

2. Ketting RF, Cochella L. Concepts and functions of small RNA pathways in C. elegans. Curr Top Dev Biol. 2021;144: 45–89. doi:10.1016/bs.ctdb.2020.08.002

3. Wu J, Yang J, Cho WC, Zheng Y. Argonaute proteins: Structural features, functions and emerging roles. Journal of Advanced Research. 2020;24: 317–324. doi:10.1016/j.jare.2020.04.017

4. Dueck A, Meister G. Assembly and function of small RNA - argonaute protein complexes. Biol Chem. 2014;395: 611–629. doi:10.1515/hsz-2014-0116

5. Meister G. Argonaute proteins: functional insights and emerging roles. Nat Rev Genet. 2013;14: 447–459. doi:10.1038/nrg3462

6. Yigit E, Batista PJ, Bei Y, Pang KM, Chen C-CG, Tolia NH, et al. Analysis of the C. elegans Argonaute Family Reveals that Distinct Argonautes Act Sequentially during RNAi. Cell. 2006;127: 747–757. doi:10.1016/j.cell.2006.09.033

7. Chen X, Wang K, Mufti FUD, Xu D, Zhu C, Huang X, et al. Germ granule compartments coordinate specialized small RNA production. Nat Commun. 2024;15: 5799. doi:10.1038/s41467-024-50027-3

8. Wedeles CJ, Wu MZ, Claycomb JM. Protection of Germline Gene Expression by the C. elegans Argonaute CSR-1. Developmental Cell. 2013;27: 664–671. doi:10.1016/j.devcel.2013.11.016

9. Seth M, Shirayama M, Gu W, Ishidate T, Conte D, Mello CC. The C. elegans CSR-1 Argonaute pathway counteracts epigenetic silencing to promote germline gene expression. Dev Cell. 2013;27: 656–663. doi:10.1016/j.devcel.2013.11.014

10. Gushchanskaia ES, Esse R, Ma Q, Lau NC, Grishok A. Interplay between small RNA pathways shapes chromatin landscapes in C. elegans. Nucleic Acids Res. 2019;47: 5603–5616. doi:10.1093/nar/gkz275

11. Conine CC, Moresco JJ, Gu W, Shirayama M, Conte D, Yates JR, et al. Argonautes Promote Male Fertility and Provide a Paternal Memory of Germline Gene Expression in C. elegans. Cell. 2013;155: 1532–1544. doi:10.1016/j.cell.2013.11.032

12. Charlesworth AG, Seroussi U, Lehrbach NJ, Renaud MS, Sundby AE, Molnar RI, et al. Two isoforms of the essential C. elegans Argonaute CSR-1 differentially regulate sperm and oocyte fertility. Nucleic Acids Research. 2021;49: 8836–8865. doi:10.1093/nar/gkab619

13. Gerson-Gurwitz A, Wang S, Sathe S, Green R, Yeo GW, Oegema K, et al. A Small RNA-Catalytic Argonaute Pathway Tunes Germline Transcript Levels to Ensure Embryonic Divisions. Cell. 2016;165: 396–409. doi:10.1016/j.cell.2016.02.040

14. Fassnacht C, Tocchini C, Kumari P, Gaidatzis D, Stadler MB, Ciosk R. The CSR-1 endogenous RNAi pathway ensures accurate transcriptional reprogramming during the oocyte-to-embryo transition in Caenorhabditis elegans. PLoS Genet. 2018;14: e1007252. doi:10.1371/journal.pgen.1007252

15. Nguyen DAH, Phillips CM. Arginine methylation promotes siRNA-binding specificity for a spermatogenesis-specific isoform of the Argonaute protein CSR-1. Nat Commun. 2021;12: 4212. doi:10.1038/s41467-021-24526-6

16. Suh N, Blelloch R. Small RNAs in early mammalian development: from gametes to gastrulation. Development. 2011;138: 1653–1661. doi:10.1242/dev.056234

17. Tabara H, Mitani S, Mochizuki M, Kohara Y, Nagata K. A small RNA system ensures accurate homologous pairing and unpaired silencing of meiotic chromosomes. The EMBO Journal. 2023;42: e105002. doi:10.15252/embj.2020105002

18. Claycomb JM, Batista PJ, Pang KM, Gu W, Vasale JJ, van Wolfswinkel JC, et al. The Argonaute CSR-1 and its 22G-RNA cofactors are required for holocentric chromosome segregation. Cell. 2009;139: 123–134. doi:10.1016/j.cell.2009.09.014

19. Wahba L, Hansen L, Fire AZ. An essential role for the piRNA pathway in regulating the ribosomal RNA pool in C. elegans. Dev Cell. 2021;56: 2295–2312.e6. doi:10.1016/j.devcel.2021.07.014

20. Barucci G, Cornes E, Singh M, Li B, Ugolini M, Samolygo A, et al. Small-RNA-mediated transgenerational silencing of histone genes impairs fertility in piRNA mutants. Nat Cell Biol. 2020;22: 235–245. doi:10.1038/s41556-020-0462-7

21. Buckley BA, Burkhart KB, Gu SG, Spracklin G, Kershner A, Fritz H, et al. A nuclear Argonaute promotes multigenerational epigenetic inheritance and germline immortality. Nature. 2012;489: 447–451. doi:10.1038/nature11352

22. Frézal L, Saglio M, Zhang G, Noble L, Richaud A, Félix M. Genome-wide association and environmental suppression of the mortal germline phenotype of wild C. elegans. EMBO reports. 2023;24: e58116. doi:10.15252/embr.202358116

23. Upton KR, Gerhardt DJ, Jesuadian JS, Richardson SR, Sánchez-Luque FJ, Bodea GO, et al. Ubiquitous L1 Mosaicism in Hippocampal Neurons. Cell. 2015;161: 228–239. doi:10.1016/j.cell.2015.03.026

24. Richardson SR, Morell S, Faulkner GJ. L1 Retrotransposons and Somatic Mosaicism in the Brain. Annual Review of Genetics. 2014;48: 1–27. doi:10.1146/annurev-genet-120213-092412

25. Perrat PN, DasGupta S, Wang J, Theurkauf W, Weng Z, Rosbash M, et al. Transposition driven genomic heterogeneity in the Drosophila brain. Science. 2013;340: 10.1126/science.1231965. doi:10.1126/science.1231965

26. Jacob-Hirsch J, Eyal E, Knisbacher BA, Roth J, Cesarkas K, Dor C, et al. Whole-genome sequencing reveals principles of brain retrotransposition in neurodevelopmental disorders. Cell Res. 2018;28: 187–203. doi:10.1038/cr.2018.8

27. Modenini G, Abondio P, Guffanti G, Boattini A, Macciardi F. Evolutionarily recent retrotransposons contribute to schizophrenia. Transl Psychiatry. 2023;13: 1–9. doi:10.1038/s41398-023-02472-9

28. Guffanti G, Gaudi S, Fallon JH, Sobell J, Potkin SG, Pato C, et al. Transposable elements and psychiatric disorders. Am J Med Genet B Neuropsychiatr Genet. 2014;165B: 201–216. doi:10.1002/ajmg.b.32225

29. Saleh A, Macia A, Muotri AR. Transposable Elements, Inflammation, and Neurological Disease. Frontiers in Neurology. 2019;10. Available: https://www.frontiersin.org/articles/10.3389/fneur.2019.00894

30. Ravel-Godreuil C, Znaidi R, Bonnifet T, Joshi RL, Fuchs J. Transposable elements as new players in neurodegenerative diseases. FEBS Letters. 2021;595: 2733–2755. doi:10.1002/1873-3468.14205

31. Burns KH. Transposable elements in cancer. Nat Rev Cancer. 2017;17: 415–424. doi:10.1038/nrc.2017.35

32. Lažetić V, Batachari LE, Russell AB, Troemel ER. Similarities in the induction of the intracellular pathogen response in Caenorhabditis elegans and the type I interferon response in mammals. Bioessays. 2023;45: e2300097. doi:10.1002/bies.202300097

33. Ashe A, Bélicard T, Le Pen J, Sarkies P, Frézal L, Lehrbach NJ, et al. A deletion polymorphism in the Caenorhabditis elegans RIG-I homolog disables viral RNA dicing and antiviral immunity. eLife. 2013;2: e00994. doi:10.7554/eLife.00994

34. Félix M-A, Wang D. Natural Viruses of Caenorhabditis Nematodes. Annu Rev Genet. 2019;53: 313–326. doi:10.1146/annurev-genet-112618-043756

35. González R, Félix M-A. *Caenorhabditis elegans* immune responses to microsporidia and viruses. Developmental & Comparative Immunology. 2024;154: 105148. doi:10.1016/j.dci.2024.105148

36. Guo X, Zhang R, Wang J, Ding S-W, Lu R. Homologous RIG-I–like helicase proteins direct RNAi-mediated antiviral immunity in C. elegans by distinct mechanisms. Proceedings of the National Academy of Sciences. 2013;110: 16085–16090. doi:10.1073/pnas.1307453110

37. Bakowski MA, Desjardins CA, Smelkinson MG, Dunbar TA, Lopez-Moyado IF, Rifkin SA, et al. Ubiquitin-Mediated Response to Microsporidia and Virus Infection in C. elegans. PLOS Pathogens. 2014;10: e1004200. doi:10.1371/journal.ppat.1004200

38. Jiang H, Chen K, Sandoval LE, Leung C, Wang D. An Evolutionarily Conserved Pathway Essential for Orsay Virus Infection of Caenorhabditis elegans. mBio. 2017;8: e00940–17. doi:10.1128/mBio.00940-17

39. Reddy KC, Dror T, Underwood RS, Osman GA, Elder CR, Desjardins CA, et al. Antagonistic paralogs control a switch between growth and pathogen resistance in C. elegans. PLoS Pathog. 2019;15: e1007528. doi:10.1371/journal.ppat.1007528

40. Sarkies P, Ashe A, Le Pen J, McKie MA, Miska EA. Competition between virus-derived and endogenous small RNAs regulates gene expression in Caenorhabditis elegans. Genome Res. 2013;23: 1258–1270. doi:10.1101/gr.153296.112

41. Troemel ER, Félix M-A, Whiteman NK, Barrière A, Ausubel FM. Microsporidia Are Natural Intracellular Parasites of the Nematode Caenorhabditis elegans. PLOS Biology. 2008;6: e309. doi:10.1371/journal.pbio.0060309

42. Félix M-A, Ashe A, Piffaretti J, Wu G, Nuez I, Bélicard T, et al. Natural and Experimental Infection of Caenorhabditis Nematodes by Novel Viruses Related to Nodaviruses. PLOS Biology. 2011;9: e1000586. doi:10.1371/journal.pbio.1000586

43. Leyva-Díaz E, Stefanakis N, Carrera I, Glenwinkel L, Wang G, Driscoll M, et al. Silencing of Repetitive DNA Is Controlled by a Member of an Unusual Caenorhabditis elegans Gene Family. Genetics. 2017;207: 529–545. doi:10.1534/genetics.117.300134

44. Sowa JN, Jiang H, Somasundaram L, Tecle E, Xu G, Wang D, et al. The Caenorhabditis elegans RIG-I Homolog DRH-1 Mediates the Intracellular Pathogen Response upon Viral Infection. J Virol. 2020;94: e01173–19. doi:10.1128/JVI.01173-19

45. Batachari LE, Dai AY, Troemel ER. Caenorhabditis elegans RIG-I-like receptor DRH-1 signals via CARDs to activate antiviral immunity in intestinal cells. Proceedings of the National Academy of Sciences of the United States of America. 2024;121: e2402126121. doi:10.1073/pnas.2402126121

46. Lažetić V, Wu F, Cohen LB, Reddy KC, Chang Y-T, Gang SS, et al. The transcription factor ZIP-1 promotes resistance to intracellular infection in Caenorhabditis elegans. Nat Commun. 2022;13: 17. doi:10.1038/s41467-021-27621-w

47. Wu X, Shi Z, Cui M, Han M, Ruvkun G. Repression of germline RNAi pathways in somatic cells by retinoblastoma pathway chromatin complexes. PLoS Genet. 2012;8: e1002542. doi:10.1371/journal.pgen.1002542

48. Willis AR, Zhao W, Sukhdeo R, Wadi L, El Jarkass HT, Claycomb JM, et al. A parental transcriptional response to microsporidia infection induces inherited immunity in offspring. Sci Adv. 2021;7: eabf3114. doi:10.1126/sciadv.abf3114

49. Fischer SEJ, Pan Q, Breen PC, Qi Y, Shi Z, Zhang C, et al. Multiple small RNA pathways regulate the silencing of repeated and foreign genes in C. elegans. Genes Dev. 2013;27: 2678–2695. doi:10.1101/gad.233254.113

50. Guang S, Bochner AF, Pavelec DM, Burkhart KB, Harding S, Lachowiec J, et al. An Argonaute transports siRNAs from the cytoplasm to the nucleus. Science (New York, NY). 2008;321: 537. doi:10.1126/science.1157647

51. Tanguy M, Véron L, Stempor P, Ahringer J, Sarkies P, Miska EA. An Alternative STAT Signaling Pathway Acts in Viral Immunity in Caenorhabditis elegans. mBio. 2017;8: e00924–17. doi:10.1128/mBio.00924-17

52. Jose AM, Kim YA, Leal-Ekman S, Hunter CP. Conserved tyrosine kinase promotes the import of silencing RNA into Caenorhabditis elegans cells. Proc Natl Acad Sci U S A. 2012;109: 14520–14525. doi:10.1073/pnas.1201153109

53. Jiang H, Leung C, Tahan S, Wang D. Entry by multiple picornaviruses is dependent on a pathway that includes TNK2, WASL, and NCK1. eLife. 2019;8: e50276. doi:10.7554/eLife.50276

54. Panek J, Gang SS, Reddy KC, Luallen RJ, Fulzele A, Bennett EJ, et al. A cullin-RING ubiquitin ligase promotes thermotolerance as part of the intracellular pathogen response in Caenorhabditis elegans. Proc Natl Acad Sci U S A. 2020;117: 7950–7960. doi:10.1073/pnas.1918417117

55. Gang SS, Grover M, Reddy KC, Raman D, Chang Y-T, Ekiert DC, et al. A pals-25 gain-of-function allele triggers systemic resistance against natural pathogens of C. elegans. PLoS Genetics. 2022;18: e1010314. doi:10.1371/journal.pgen.1010314

56. Chen K, Franz CJ, Jiang H, Jiang Y, Wang D. An evolutionarily conserved transcriptional response to viral infection in Caenorhabditis nematodes. BMC Genomics. 2017;18: 303. doi:10.1186/s12864-017-3689-3

57. Engelmann I, Griffon A, Tichit L, Montañana-Sanchis F, Wang G, Reinke V, et al. A Comprehensive Analysis of Gene Expression Changes Provoked by Bacterial and Fungal Infection in C. elegans. PLOS ONE. 2011;6: e19055. doi:10.1371/journal.pone.0019055

58. Schott DH, Cureton DK, Whelan SP, Hunter CP. An antiviral role for the RNA interference machinery in Caenorhabditis elegans. Proceedings of the National Academy of Sciences. 2005;102: 18420–18424. doi:10.1073/pnas.0507123102

59. Niescierowicz K, Pryszcz L, Navarrete C, Tralle E, Sulej A, Abu Nahia K, et al. Adar-mediated A-to-I editing is required for embryonic patterning and innate immune response regulation in zebrafish. Nat Commun. 2022;13: 5520. doi:10.1038/s41467-022-33260-6

60. Liddicoat BJ, Piskol R, Chalk AM, Ramaswami G, Higuchi M, Hartner JC, et al. RNA editing by ADAR1 prevents MDA5 sensing of endogenous dsRNA as nonself. Science. 2015;349: 1115–1120. doi:10.1126/science.aac7049

61. Mannion NM, Greenwood SM, Young R, Cox S, Brindle J, Read D, et al. The RNA-editing enzyme ADAR1 controls innate immune responses to RNA. Cell Rep. 2014;9: 1482–1494. doi:10.1016/j.celrep.2014.10.041

62. Reich DP, Tyc KM, Bass BL. C. elegans ADARs antagonize silencing of cellular dsRNAs by the antiviral RNAi pathway. Genes Dev. 2018;32: 271–282. doi:10.1101/gad.310672.117

63. Eisenberg E, Levanon EY. A-to-I RNA editing — immune protector and transcriptome diversifier. Nat Rev Genet. 2018;19: 473–490. doi:10.1038/s41576-018-0006-1

64. Ganem NS, Ben-Asher N, Manning AC, Deffit SN, Washburn MC, Wheeler EC, et al. Disruption in A-to-I Editing Levels Affects C. elegans Development More Than a Complete Lack of Editing. Cell Rep. 2019;27: 1244–1253.e4. doi:10.1016/j.celrep.2019.03.095

65. Reich DP, Bass BL. Inverted repeat structures are associated with essential and highly expressed genes on C. elegans autosome distal arms. RNA. 2018;24: 1634–1646. doi:10.1261/rna.067405.118

66. Fischer SEJ, Montgomery TA, Zhang C, Fahlgren N, Breen PC, Hwang A, et al. The ERI-6/7 helicase acts at the first stage of an siRNA amplification pathway that targets recent gene duplications. PLoS Genet. 2011;7: e1002369. doi:10.1371/journal.pgen.1002369

67. Lev I, Gingold H, Rechavi O. H3K9me3 is required for inheritance of small RNAs that target a unique subset of newly evolved genes. Elife. 2019;8: e40448. doi:10.7554/eLife.40448

68. Newman MA, Ji F, Fischer SEJ, Anselmo A, Sadreyev RI, Ruvkun G. The surveillance of pre-mRNA splicing is an early step in C. elegans RNAi of endogenous genes. Genes Dev. 2018;32: 670–681. doi:10.1101/gad.311514.118

69. Blango MG, Bass BL. Identification of the long, edited dsRNAome of LPS-stimulated immune cells. Genome Res. 2016;26: 852–862. doi:10.1101/gr.203992.116

70. Simmer F, Tijsterman M, Parrish S, Koushika SP, Nonet ML, Fire A, et al. Loss of the putative RNA-directed RNA polymerase RRF-3 makes C. elegans hypersensitive to RNAi. Curr Biol. 2002;12: 1317–1319. doi:10.1016/s0960-9822(02)01041-2

71. Reddy KC, Dror T, Sowa JN, Panek J, Chen K, Lim ES, et al. An Intracellular Pathogen Response Pathway Promotes Proteostasis in C. elegans. Curr Biol. 2017;27: 3544–3553.e5. doi:10.1016/j.cub.2017.10.009

72. Cook DE, Zdraljevic S, Roberts JP, Andersen EC. CeNDR, the Caenorhabditis elegans natural diversity resource. Nucleic Acids Res. 2017;45: D650–D657. doi:10.1093/nar/gkw893

73. Widmayer SJ, Evans KS, Zdraljevic S, Andersen EC. Evaluating the power and limitations of genome-wide association studies in Caenorhabditis elegans. G3: Genes|Genomes|Genetics. 2022;12: jkac114. doi:10.1093/g3journal/jkac114

74. Lee D, Zdraljevic S, Stevens L, Wang Y, Tanny RE, Crombie TA, et al. Balancing selection maintains hyper-divergent haplotypes in C. elegans. Nature ecology & evolution. 2021;5: 794. doi:10.1038/s41559-021-01435-x

75. Zhang G, Roberto NM, Lee D, Hahnel SR, Andersen EC. The impact of species-wide gene expression variation on Caenorhabditis elegans complex traits. Nat Commun. 2022;13: 3462. doi:10.1038/s41467-022-31208-4

76. Zhou H, He Y, Huang Y, Li R, Zhang H, Xia X, et al. Comprehensive analysis of prognostic value, immune implication and biological function of CPNE1 in clear cell renal cell carcinoma. Front Cell Dev Biol. 2023;11. doi:10.3389/fcell.2023.1157269

77. Ramsey CS, Yeung F, Stoddard PB, Li D, Creutz CE, Mayo MW. Copine-I represses NF-κB transcription by endoproteolysis of p65. Oncogene. 2008;27: 3516–3526. doi:10.1038/sj.onc.1211030

78. Yin X, Zou B, Hong X, Gao M, Yang W, Zhong X, et al. Rice copine genes Os1 and Os3 function as suppressors of broad-spectrum disease resistance. Plant Biotechnology Journal. 2018;16: 1476–1487. doi:10.1111/pbi.12890

79. Zou B, Ding Y, Liu H, Hua J. Silencing of copine genes confers common wheat enhanced resistance to powdery mildew. Molecular Plant Pathology. 2018;19: 1343–1352. doi:10.1111/mpp.12617

80. Duchaine TF, Wohlschlegel JA, Kennedy S, Bei Y, Conte D, Pang K, et al. Functional Proteomics Reveals the Biochemical Niche of C. elegans DCR-1 in Multiple Small-RNA-Mediated Pathways. Cell. 2006;124: 343–354. doi:10.1016/j.cell.2005.11.036

81. Warner A, Xiong G, Qadota H, Rogalski T, Vogl AW, Moerman DG, et al. CPNA-1, a copine domain protein, is located at integrin adhesion sites and is required for myofilament stability in Caenorhabditis elegans. Mol Biol Cell. 2013;24: 601–616. doi:10.1091/mbc.E12-06-0478

82. Raman D, Wernet N, Gang S, Troemel E. PALS-14 promotes resistance to Nematocida parisii infection in Caenorhabditis elegans. microPublication Biology. 2024 [cited 23 Nov 2024]. doi:10.17912/micropub.biology.001325

83. van Sluijs L, Bosman KJ, Pankok F, Blokhina T, Wilten JIHA, te Molder DM, et al. Balancing Selection of the Intracellular Pathogen Response in Natural Caenorhabditis elegans Populations. Front Cell Infect Microbiol. 2022;11. doi:10.3389/fcimb.2021.758331

84. Balla KM, Andersen EC, Kruglyak L, Troemel ER. A Wild C. Elegans Strain Has Enhanced Epithelial Immunity to a Natural Microsporidian Parasite. PLoS Pathogens. 2015;11: e1004583. doi:10.1371/journal.ppat.1004583

85. Spichal M, Heestand B, Billmyre KK, Frenk S, Mello CC, Ahmed S. Germ granule dysfunction is a hallmark and mirror of Piwi mutant sterility. Nat Commun. 2021;12: 1420. doi:10.1038/s41467-021-21635-0

86. Wang D, Zhang Z, O’Loughlin E, Lee T, Houel S, O’Carroll D, et al. Quantitative functions of Argonaute proteins in mammalian development. Genes Dev. 2012;26: 693–704. doi:10.1101/gad.182758.111

87. Pal A, Vasudevan V, Houle F, Lantin M, Maniates KA, Huberdeau MQ, et al. Defining the contribution of microRNA-specific Argonautes with slicer capability in animals. Nucleic Acids Res. 2024;52: 5002–5015. doi:10.1093/nar/gkae173

88. Bouasker S, Simard MJ. The slicing activity of miRNA-specific Argonautes is essential for the miRNA pathway in C. elegans. Nucleic Acids Research. 2012;40: 10452. doi:10.1093/nar/gks748

89. Lažetić V, Blanchard MJ, Bui T, Troemel ER. Multiple pals gene modules control a balance between immunity and development in Caenorhabditis elegans. PLoS Pathog. 2023;19: e1011120. doi:10.1371/journal.ppat.1011120

90. Martinez JC, Morandini F, Fitzgibbons L, Sieczkiewicz N, Bae SJ, Meadow ME, et al. cGAS deficient mice display premature aging associated with de-repression of LINE1 elements and inflammation. bioRxiv; 2024. p. 2024.10.10.617645. doi:10.1101/2024.10.10.617645

91. Brenner S. The genetics of Caenorhabditis elegans - PubMed. 1974;77. Available: https://pubmed.ncbi.nlm.nih.gov/4366476/

92. Kamath RS, Ahringer J. Genome-wide RNAi screening in Caenorhabditis elegans. Methods. 2003;30: 313–321. doi:10.1016/s1046-2023(03)00050-1

93. Rual J-F, Ceron J, Koreth J, Hao T, Nicot A-S, Hirozane-Kishikawa T, et al. Toward Improving Caenorhabditis elegans Phenome Mapping With an ORFeome-Based RNAi Library. Genome Res. 2004;14: 2162–2168. doi:10.1101/gr.2505604

94. Teichman G, Cohen D, Ganon O, Dunsky N, Shani S, Gingold H, et al. RNAlysis: analyze your RNA sequencing data without writing a single line of code. BMC Biology. 2023;21: 74. doi:10.1186/s12915-023-01574-6

95. Martin M. Cutadapt removes adapter sequences from high-throughput sequencing reads. EMBnet.journal. 2011;17: 10–12. doi:10.14806/ej.17.1.200

96. Shahid S, Axtell MJ. Identification and annotation of small RNA genes using ShortStack. Methods. 2014;67: 20–27. doi:10.1016/j.ymeth.2013.10.004

97. Liao Y, Smyth GK, Shi W. featureCounts: an efficient general purpose program for assigning sequence reads to genomic features. Bioinformatics. 2014;30: 923–930. doi:10.1093/bioinformatics/btt656

98. Bray NL, Pimentel H, Melsted P, Pachter L. Near-optimal probabilistic RNA-seq quantification. Nat Biotechnol. 2016;34: 525–527. doi:10.1038/nbt.3519

99. Love MI, Huber W, Anders S. Moderated estimation of fold change and dispersion for RNA-seq data with DESeq2. Genome Biology. 2014;15: 550. doi:10.1186/s13059-014-0550-8

100. Ghanta KS, Mello CC. Melting dsDNA Donor Molecules Greatly Improves Precision Genome Editing in Caenorhabditis elegans. Genetics. 2020;216: 643–650. doi:10.1534/genetics.120.303564

101. Frézal L, Demoinet E, Braendle C, Miska E, Félix M-A. Natural Genetic Variation in a Multigenerational Phenotype in C. elegans. Curr Biol. 2018;28: 2588–2596.e8. doi:10.1016/j.cub.2018.05.091

102. Batachari LE, Sarmiento MB, Wernet N, Troemel ER. Orsay Virus Infection in Caenorhabditis elegans. Curr Protoc. 2024;4: e1098. doi:10.1002/cpz1.1098

103. Ghaddar A, Armingol E, Huynh C, Gevirtzman L, Lewis NE, Waterston R, et al. Whole-body gene expression atlas of an adult metazoan. Science Advances. 2023;9: eadg0506. doi:10.1126/sciadv.adg0506

104. Wan G, Fields BD, Spracklin G, Shukla A, Phillips CM, Kennedy S. Spatiotemporal regulation of liquid-like condensates in epigenetic inheritance. Nature. 2018;557: 679–683. doi:10.1038/s41586-018-0132-0

